# White matter structural bases for predictive tapping synchronization

**DOI:** 10.1101/2022.09.05.506691

**Authors:** Pamela García-Saldivar, Cynthia de León, Luis Concha, Hugo Merchant

**Author notes:** Corresponding authors. *Email addresses.

## Abstract

We determined the intersubject association between rhythmic entrainment abilities of human subjects during a synchronization continuation tapping task (SCT) and the macro and microstructural properties of their superficial (SWM) and deep (dWM) white matter. Diffusion-weighted images were obtained from 32 subjects who also performed the SCT with auditory or visual metronomes and five tempos ranging from 550 to 950 ms. We developed a method to determine the fiber density of U-fibers running tangentially to the cortex. Notably, the right audiomotor system showed individual differences in the density of U-fibers that were correlated with the degree of predictive entrainment across subjects. These correlations were selective for the synchronization epoch with auditory metronomes and were specific for tempos around 1.5 Hz. In addition, there was a significant association between predictive rhythmic entrainment and the density and bundle diameter of the corpus callosum (CC), forming a chronotopic map where behavioural correlations of short and long intervals were found with the anterior and posterior portions of the CC. Finally, the fiber bundle cross-section of the arcuate fasciculus, the CC, and the Superior Longitudinal Fasciculus showed a significant correlation with the mean asynchronies of the auditory SCT. These findings suggest that the structural properties of the SWM and dWM in the audiomotor system support the predictive abilities of subjects during rhythmic tapping, where the density of cortical U-fibers are linked to the preferred tapping tempo, and the bundle properties of CC define an interval selective topography that has an anterior posterior gradient.

## 1. Introduction

Moving in synchrony with a beat is a basic and generalized human ability that can reach sophisticated levels in professional percussionist (Honing et al., 2015; Merchant et al., 2015a). Indeed, humans are extremely sensitive to auditory regularities, can entrain to auditory beats across a wide range of tempos, and can use timed movements of different body parts (such as finger or foot taps, or body sway) to keep the beat (Mendoza & Merchant, 2014; Repp & Su, 2013). A classical task used to study rhythmic entrainment is the synchronization-continuation task (SCT), where subjects first entrain their tapping to an isochronous metronome and then continue tapping without the periodic stimulus using an internal clock (Merchant et al., 2008a,b; Wing, 2002; Wing & Kristofferson, 1973). In this type of tasks humans show negative asynchronies, namely they tap a few milliseconds before the metronome, supporting the notion that entrainment depends on a predictive internal pulse (Lenc et al., 2021; Repp & Su, 2013; Zarco et al., 2009). Neurophysiological and functional imaging studies have shown that the internal beat representation during SCT rests in the motor system including the basal ganglia and the supplementary motor regions (Bartolo et al., 2014; Merchant et al., 2013a, 2015a; Merchant & Averbeck, 2017; Rao et al., 2001; Sánchez-Moncada et al., 2020). These areas produce a regenerating rhythmic signal (Crowe et al., 2014; Gámez et al., 2019; Merchant et al., 2015b) that dynamically interacts with the auditory areas, creating audiomotor loops where the motor prediction of the pulse is flexibly compared and adjusted depending on the changes in the input stream of rhythmic stimuli (Comstock et al., 2018; Merchant & Honing, 2014; Patel & Iversen, 2014). In addition, the supplementary motor areas possess neurons that are tuned to the tempo duration of tapping (Merchant et al., 2014, 2013b) giving rise to interval specific circuits that define a chronotopic map, with short preferred intervals in the anterior portion and long preferred intervals in the posterior portion of the medial premotor areas (Protopapa et al., 2019).

The existence of neural circuits with preferred intervals conforming chronotopic maps goes in line with the notion of a spontaneous rhythmic tempo, an interval that is produced naturally when asked to tap in the absence of external cues (McAuley et al., 2006; Zamm et al., 2018). This spontaneous or preferred tempo is around 600 to 750 ms in human adults (Fraisse, 1963, 1978) being faster in early childhood and slower in elderly individuals (McAuley et al., 2006). Thus, the biological intrinsic periodicity may depend on a biased distribution of preferred tempos towards 1.5 Hz in the interval-tuned neurons of the motor system (Bartolo et al., 2014; Bartolo & Merchant, 2009; Merchant et al., 2013b).

Although rhythmic entrainment is prevalent across all human cultures and is a natural behaviour for social interaction (Nettle, 2000), there are wide individual differences in the precision, accuracy and predictability (measured with the asynchronies that correspond to the time between the stimulus and the response) of movement synchronization. There are subjects who, despite lacking musical training, spontaneously synchronize to rhythmic stimuli ranging from strictly periodic metronomes to complex musical pieces, with performance comparable to trained musicians (Scheurich et al., 2018; Tranchant et al., 2016). In contrast, there are also poor synchronizers (around ten percent of the population) that show low accuracy and large asynchronies to isochronous metronomes and musical excerpts (Phillips-Silver et al., 2011; Sowiński & Dalla Bella, 2013; Tranchant et al., 2016). Furthermore, non-musicians synchronize less flexibly and less precisely across tempi than musicians (Scheurich et al., 2018). Hence, both genetic and learning factors generate human populations with a wide variety of beat entrainment abilities. Notably, functional imaging has revealed that individual differences in beat perception depend on activation differences in SMA and the posterior auditory cortex (Grahn & McAuley, 2009). In addition, the putamen, SMA and auditory cortex show larger functional connectivity during rhythm perception, with larger modulation for musicians than non-musicians (Grahn & Rowe, 2009, 2013). Consequently, the magnitude of the anatomofunctional association between the auditory and motor control areas of the cerebral cortex seem to covary with the individual difference to perceive and entrain to simple regular beats. Such brain networks rely on the structural connectivity provided by white matter. Yet, little is known about whether the macro and microstructural properties in the superficial and deep white matter can explain the individual variations in rhythmic entrainment performance (Assaneo et al., 2019; Steele et al., 2013). We hypothesized that the properties in the microstructure of fibers running superficially to the cortical surface and of the deep tracts of the audiomotor system account for the individual differences of the predictive abilities during tapping synchronization.

To test this hypothesis, we acquired diffusion-weighted images (DWI) from 32 subjects that previously performed a SCT using flashing visual or auditory tones as metronomes in the ranges of hundreds of milliseconds. SCT rhythmic performance across durations (550,650,750,850 or 950 ms) and modalities (auditory and visual) was characterized using the absolute asynchronies, the autocorrelation of the inter-tap interval time series during the synchronization epoch, the constant error, and the temporal variability during both synchronization and continuation epochs. These parameters measure the prediction, error correction, accuracy, and precision of the rhythmic tapping of the subjects, respectively (Figure 2A). On the other hand, we evaluated deep white matter fascicles using a fixel-based approach (Dhollander et al., 2021), and developed two metrics of superficial white matter, namely fiber density corresponding to fibers entering/exiting the cortex and U-fibers running tangentially to the cortex. Crucially, we used a wide range of tempos in the SCT in order to test the existence of both a structural foundation for the preferred interval and an interval selective topographical arrangement in the behavioural and structural associations. Widespread correlations in the right audiomotor circuit were found between the tangential U-fibers density and the predictive abilities of subjects during synchronization epoch of the auditory condition for the intervals of 650 and 750ms. The interval specificity in these associations suggest that the preferred tempo for rhythmic entrainment has its origins in the structural properties of the U-fibers running superficially across the audiomotor circuit. In addition, there was a significant association between predictive rhythmic entrainment in the auditory condition and the density and bundle diameter of the corpus callosum, forming a chronotopic map with an anterior-posterior gradient, similar to the topography of interval tuned clusters observed with functional imaging. Crucially, the anatomobehavioural associations were negative, indicating that subjects with good predictive abilities and small asynchronies showed large superficial and deep apparent fiber densities, while subjects with large asynchronies showed low fiber densities.

## 2. Results

### 2.1. Rhythmic behaviour

Thirty-two subjects performed a modified version of the classical synchronization continuation task (SCT) that included the following three epochs: beat perception, synchronization, and continuation (Figure 1). This task starts with the active perception of the isochronous beat defined by visual stimuli alternating between a right and left position, followed by tapping synchronization to the alternating stimuli and the internally driven tapping continuation to the right or left targets without the metronome. The subjects also performed an auditory version of the SCT (Figure 1, see Methods).

**Figure 1:**
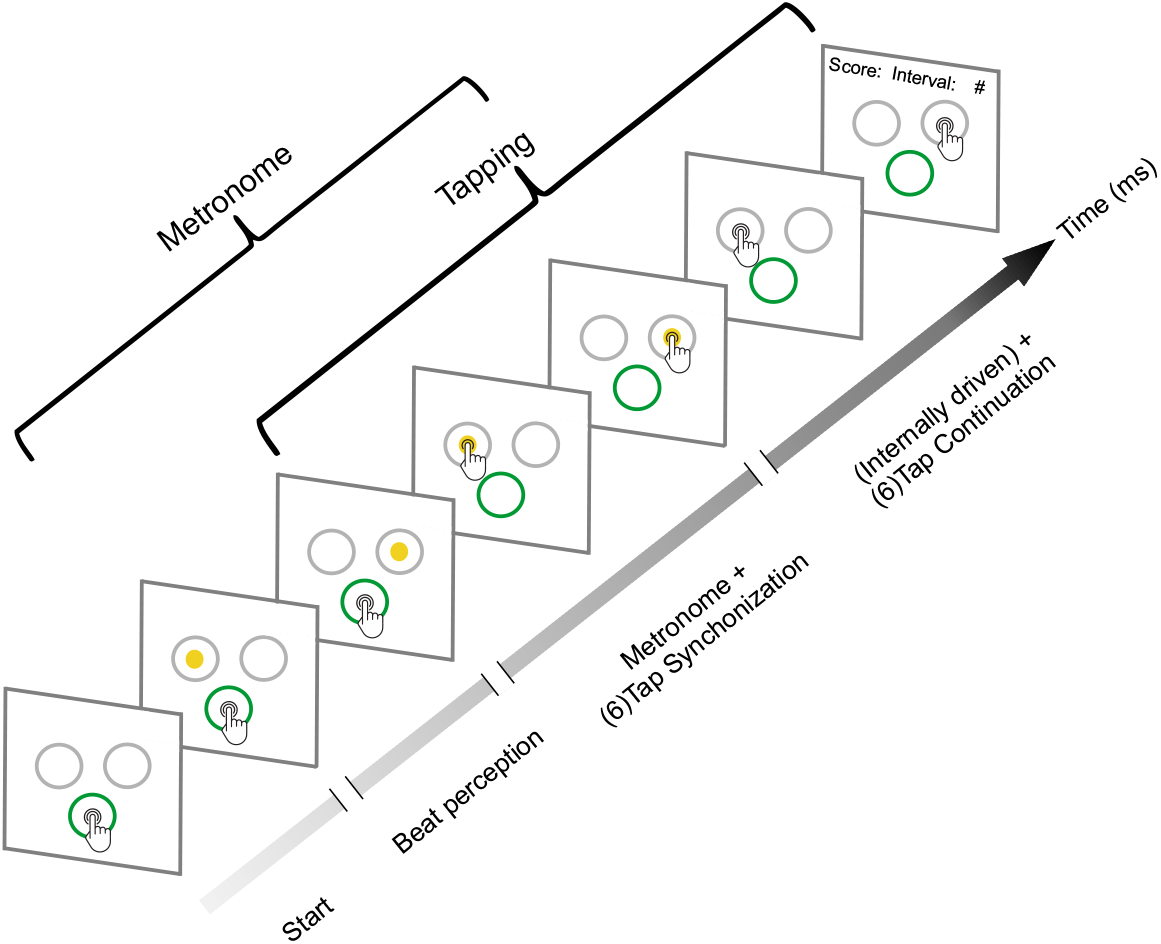
Synchronization-Continuation Task (SCT). Initially, the subjects placed their finger at the central-bottom target of a touch screen to start the trial and maintained the finger in this position while attending to a sequence of two to six right/left alternating visual stimuli with a constant interstimulus interval (target duration, perception epoch). Then, once they considered to have extracted the beat, subjects tapped the touch screen on the left or right target in synchrony with the alternating visual metronome during six intervals (synchronization epoch). Finally, they continued taping on the right or left targets for another six intervals without the metronome (continuation epoch). The mean produced and the target interval were displayed at the end of each trial as feedback. Subjects also performed an auditory metronome version of the SCT, where the metronome consisted of 500 Hz tones that were presented on the right or left side of a headphone. Five target durations (550,650,750,850 or 950 ms) were presented pseudorandomly, with the visual and auditory conditions interleaved between subjects.

Absolute asynchronies correspond to the time difference between each stimulus and response pair and are a measure of rhythmic prediction. Hence, this parameter can only be measured during the synchronization epoch of SCT. A repeated-measures ANOVA on absolute asynchronies with metronome modality (auditory and visual) and instructed interval duration as within-subject factor, showed significant main effects for duration (F(4,124) = 36.88 P < 0.0001) and modality (F(1,31) = 20 P < 0.0001), as well as a significant duration x modality interaction (F(4,124) = 32.3 P < 0.0001). Tukey’s honest significant difference (HSD) post hoc test showed significantly larger asynchronies for the visual than auditory modality across all durations (P < 0.0001 for 550,650,750 and 850 ms; P = 0.006 for 950 ms). These results confirm the preponderance of the auditory over the visual modality to guide rhythmic temporal prediction (Comstock et al., 2018; Gámez et al., 2018; Merchant et al., 2015b). In addition, we computed the intersubject correlation matrix on the absolute asynchronies across instructed intervals and we found a significant correlation between 650 and 750 ms for both the auditory and visual conditions (r=0.67, p=.000026; r=0.69, p=.000011, respectively) (Extended Data 1). This finding suggests the existence of a shared mechanism for rhythmic temporal prediction in these intermediate intervals.

A negative lag 1 autocorrelation of the produced intervals during the synchronization epoch indicates the involvement of an error correction mechanism that maintains tap synchronization with the metronome, since a longer produced interval tends to be followed by a shorter interval, while a shorter interval tends to be followed by a longer produced duration (Iversen et al., 2015; Repp & Su, 2013). The corresponding repeated-measures ANOVA on autocorrelation of the inter-tap interval time series (Figure 2C) showed no significant differences between modalities (F(1,31) = 1.6 P = 0.21), intervals (F(4,124) = 2.05 P = 0.09) or their interaction (F(4,124) = 2.34 P = 0.06). Thus, lag 1 autocorrelation across trials was negative and similar across modalities and target durations supporting the notion of a robust and amodal error correction mechanism during the SCT.

**Figure 2:**
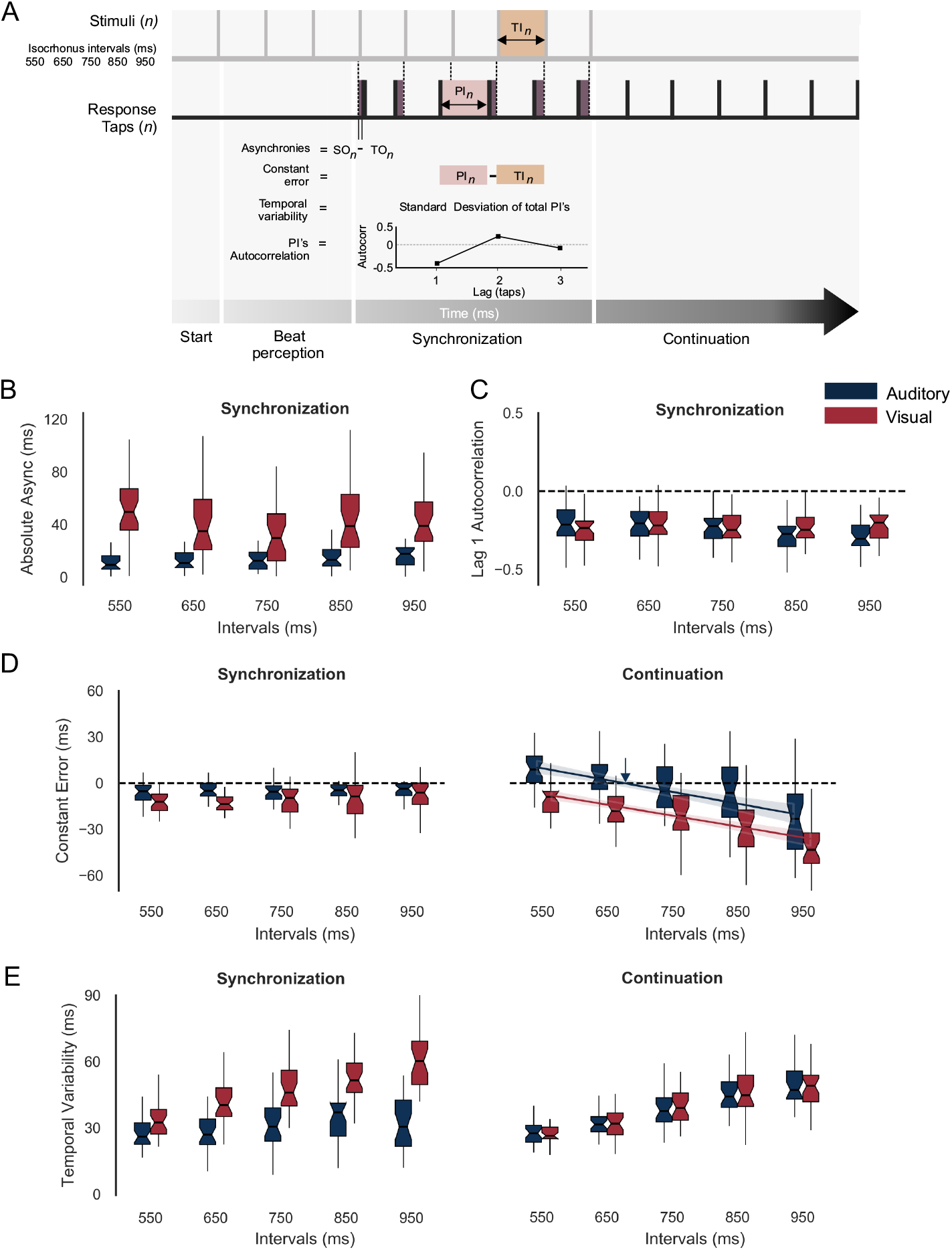
A. Parameters of rhythmic performance during SCT. Asynchronies correspond to the time difference between stimulus (SO) and tap onset (TO) across the n intervals of the synchronization epoch. The constant error is the difference between produced (PI) and target intervals (TI), and the temporal variability is the standard deviation of the PI. Finally, the autocorrelation of the PI during synchronization is computed and the lag 1 autocorrelation: a negative value indicates that the subject is using an error correction mechanism (see the text). B and C. Absolute asynchronies and lag 1 autocorrelation for each instructed interval and metronome modality (Auditory: blue, Visual: red) during the synchronization epoch. D and E. Constant error and temporal variability as a function of target interval for both metronome modalities and the two tapping epochs of the SCT. The coloured lines in D for the continuation correspond to the linear fit between the constant error and the target interval; the indifference interval corresponds to 680ms (blue vertical arrow) for the auditory condition.

Constant error is the difference between the produced and the target intervals and is a measure of timing accuracy during the synchronization and continuation epochs. A repeated-measures ANOVA on constant error with modality, target duration and task epoch as within-subject factors, revealed statistically significant main effects for modality (F(1,31) = 46.05, P < 0.0001), target duration (F(4,124) = 29.15, P < 0.0001) and epoch (F(1,31) = 6.44, P = 0.01), as well significant interactions between: modality x interval (F(4,124) = 3.5, P = 0.008), epoch x interval (F(4,124) = 48.33, P < 0.0001), and epoch x modality (F(1,31) = 23.35, P < 0.0001). The post hoc Tukey HSD showed significant no differences for target duration in constant error during synchronization for both modalities, with accurate timing close to zero. In contrast, during continuation, the same post hoc test revealed significant differences between distant intervals within the auditory and visual modalities (Figure 2D). For the continuation epoch, the constant error followed the bias effect, with overestimation for short and underestimation for long durations, especially for the auditory condition (Jazayeri & Shadlen, 2010; Pérez & Merchant, 2018). Indeed, the indifference interval, that corresponds to the interval associated with zero constant error, was 654ms for the auditory (Figure 2D, blue arrow) and 420 ms for the visual condition. This finding suggests that our subjects had a clear preferred interval in the auditory condition that is close to the reported 2Hz in the literature (Zamm et al., 2016). Finally, temporal variability was defined as the standard deviation of the produced intervals and is a metric of timing precision. The same repeated-measures ANOVA on temporal variability showed significant main effects for target duration (F(4,124) = 110, P < 0.0001), and modality (F(1,31) = 58.06, P < 0.0001), but no significant main effect for task epoch (F(1,31)=0.94, P =0.33). In addition, significant effects were revealed for the following interactions: epoch-interval (F(4,124) = 3.42, P = 0.01), epoch-modality (F(1,31) = 70.68, P < 0.0001), interval-modality (F(4,124) = 11.34, P = 0.0001), and epoch-modality-interval (F(4,124) = 14.74, P = 0.0001). The HSD post hoc tests showed significantly larger temporal variability in the visual than the auditory metronome (for the intervals 650, 750, 850, and 950: P < 0.0001) during the synchronization but not the continuation epoch, confirming the larger precision for auditory metronomes, especially during synchronization (Gámez et al., 2018; Repp & Penel, 2004).

### 2.2. Analysis of white matter

Before the task performance session, participants were scanned in a 3 T Philips Achieva TX MR scanner with a 32-channel head coil. T1-weighted volumes and diffusion-weighted images (DWI) were obtained (see Methods). For each subject, the gray/white matter interface was identified using a surface mesh (Garcia-Saldivar et al., 2021). The apparent fiber density (AFD) (Raffelt et al., 2012), derived from the DWI using constrained spherical deconvolution (CSD) (Tournier et al., 2004), was sampled at each vertex of this mesh. To evaluate white matter properties at different depths with respect to the cortical mantle, we created synthetic trajectories that organically extend from each vertex at the grey/white matter interface towards the ventricles and sampled diffusion metrics along these trajectories every 0.5 mm (Figure 3A). Leveraging the ability of CSD to disentangle crossing fiber populations, AFD (Figure 3B) was evaluated separately for those fibers that enter/exit the cortex and are, therefore, parallel to the virtual trajectories (*_par_AFD*), and those that extend tangentially to the cortex, perpendicular to the virtual trajectories (*_tan_AFD*). The latter corresponds to short-range cortico-cortical connections through U-fibers (Schüz & Braitenberg, 2002).

**Figure 3:**
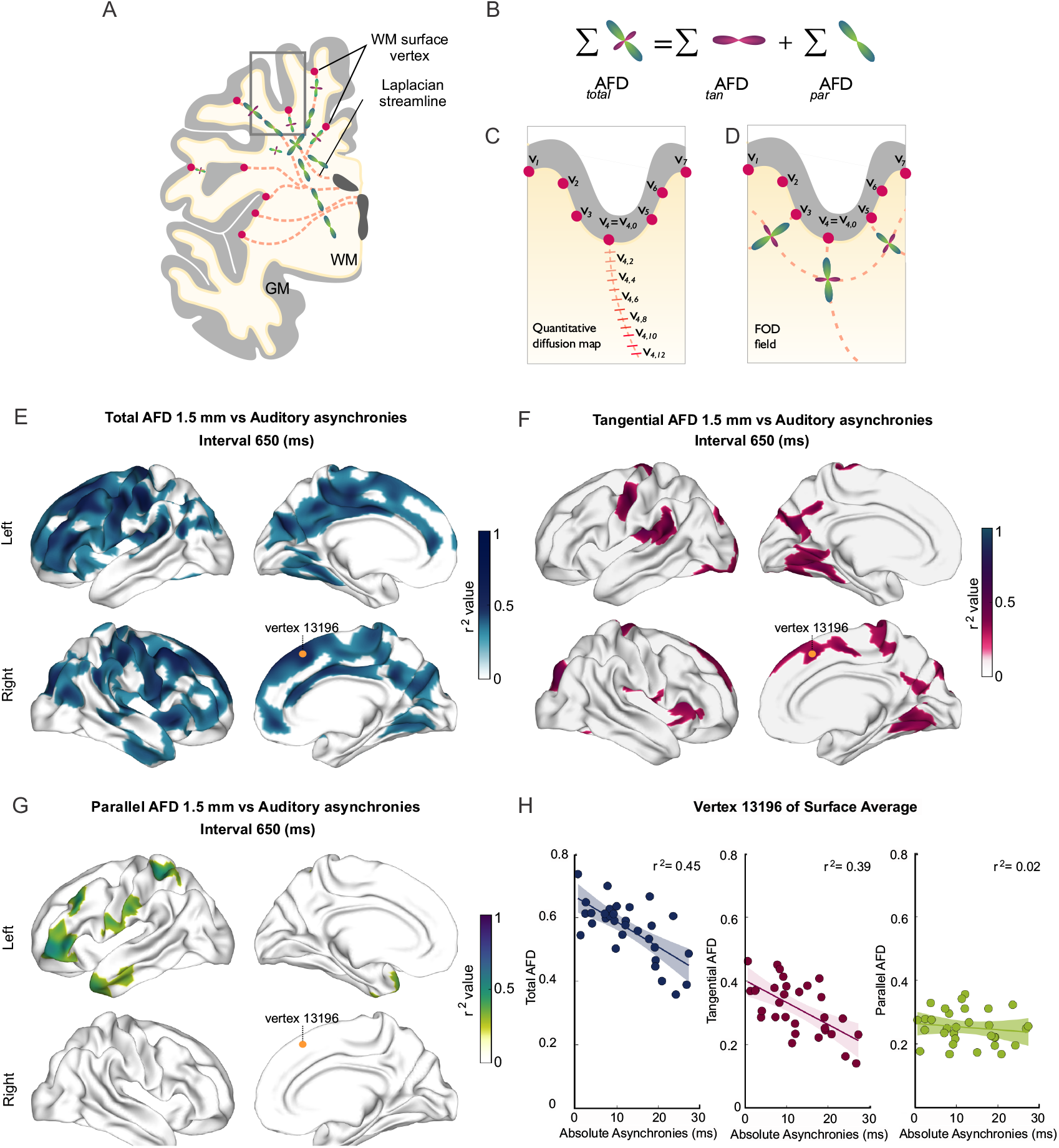
SWM analysis. Apparent fiber density (AFD) of superficial white matter was systematically sampled along synthetic streamlines that extend from each vertex of the gray/white matter interface surface towards the ventricles following a Laplacian field (A, C). The integral of all fiber orientation distribution functions (FOD) corresponds to the total apparent fiber density (*_total_AFD*), further separated (B) into fiber densities corresponding to fibers entering/exiting the cortex parallel to the Laplacian streamlines (*_par_AFD*), or U-fibers running tangentially to the cortex (*_tan_AFD*) (D). (E) There are widespread significant correlations between the subjects’ asynchronies during the SCT with an auditory metronome and *_total_AFD*, shown here for the 650 ms interval (E). (F) Large areas within the frontal, parietal and occipital lobes showed significant correlations between behaviour and *_tan_AFD*, while only restricted frontal and temporal regions showed correlation between *_par_AFD* and the asynchronies (G). Panel H shows the correlation across the 32 subjects between the three AFD metrics and asynchronies for one exemplary vertex (yellow dot in E, F, and G).

### 2.3. Correlations between behaviour and Superficial White Matter (SWM)

Next, we examined the association between behavioural performance and microstructural properties of superficial white matter, inferred from diffusion-weighted magnetic resonance images. The surface-based analysis of SWM (SBA, see Methods) was performed to determine the possible association between the different metrics of rhythmic timing performance and the metrics of the SWM sampled at five depths with respect to the grey/white matter interface (0, 0.5, 1, 1.5 and 2 mm). This analysis showed the existence of negative correlations between the auditory absolute asynchronies and values of the AFD maps in the five depths sampled (Figure 3 EFG; also see Extended Data 2). Thus, subjects with auditory asynchronies closer to zero, and hence with larger predictive abilities, had significantly higher values AFD and *_tan_AFD* than subjects with less predictive performance. Notably, these significant associations were observed mainly for the auditory asynchronies of the 650 and 750 ms intervals and the values of *_total_AFD* and *_tan_AFD* (see Extended Data 3, 4 and 5). No significant correlations were found between the three AFD maps and the constant error, temporal variability, and lag 1 autocorrelation for the auditory condition. Furthermore, no correlations were observed between the three metrics of the SWM and all the behavioural parameters for the visual condition (see Extended Data 2). We did not find significant correlations between the parameters of SCT rhythmic performance and the *_par_AFD*, except for few clusters with low correlation coefficients for the auditory asynchronies of the intervals of 550, 650 and 750 ms (Extended Data 2 and Figure 3H). Indeed, the level of association between predictive behaviour and the AFD maps was larger for *_tan_AFD* than *_par_AFD* (Figure 3H; compare Figure 5 with Extended Data 8). The average thickness of the U-fibre system is around 1.5 mm (Schüz & Braitenberg, 2002), and following analyses focus on this depth. Figure 4 shows the correlation coefficient values between the auditory asynchronies and *_tan_AFD* at 1.5mm below the GM/WM interface across all the tested tempos. After correction for multiple comparisons (*p_FWE_* < 0.001 and *p_cluster_* < 0.001), only intervals of 650, 750 and 850ms showed a significant association between the behavioural and structural parameter. Indeed, nine (with 2232 vertices), twelve (with 2827 vertices), and two (with 374 vertices) clusters showed significant correlations between auditory asynchronies and *_tan_AFD* for the 650, 750 and 850ms intervals, respectively.

**Figure 4:**
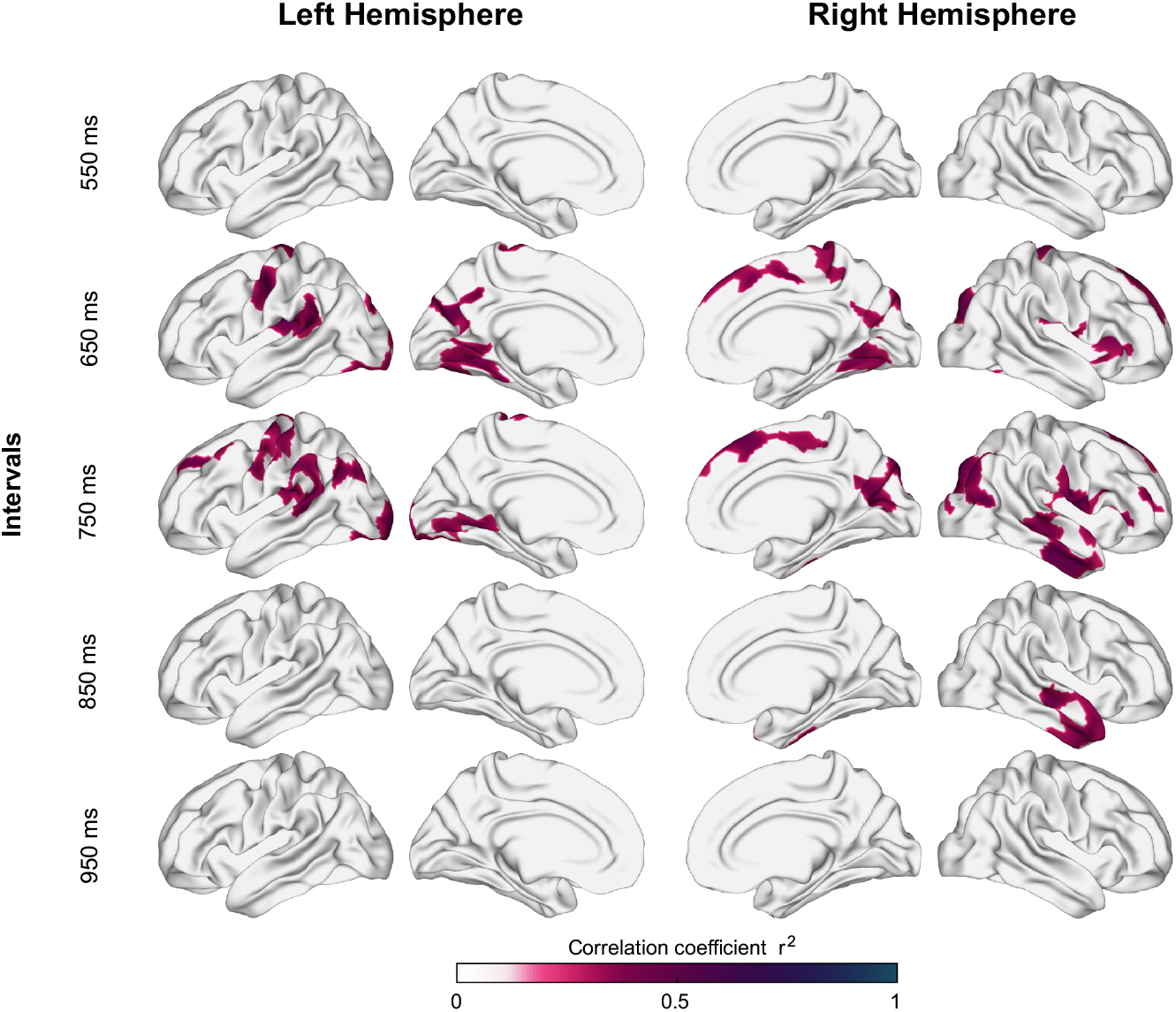
Correlation (r^2^) between auditory asynchronies and tangential superficial white matter apparent fiber density for each vertex (sampled at 1.5 mm below the gray/white matter interface) across all tested intervals in the SCT for both hemispheres. Significant correlations were localized in large clusters within motor, auditory, and visual areas, particularly for 650 and 750 ms intervals.

**Figure 5:**
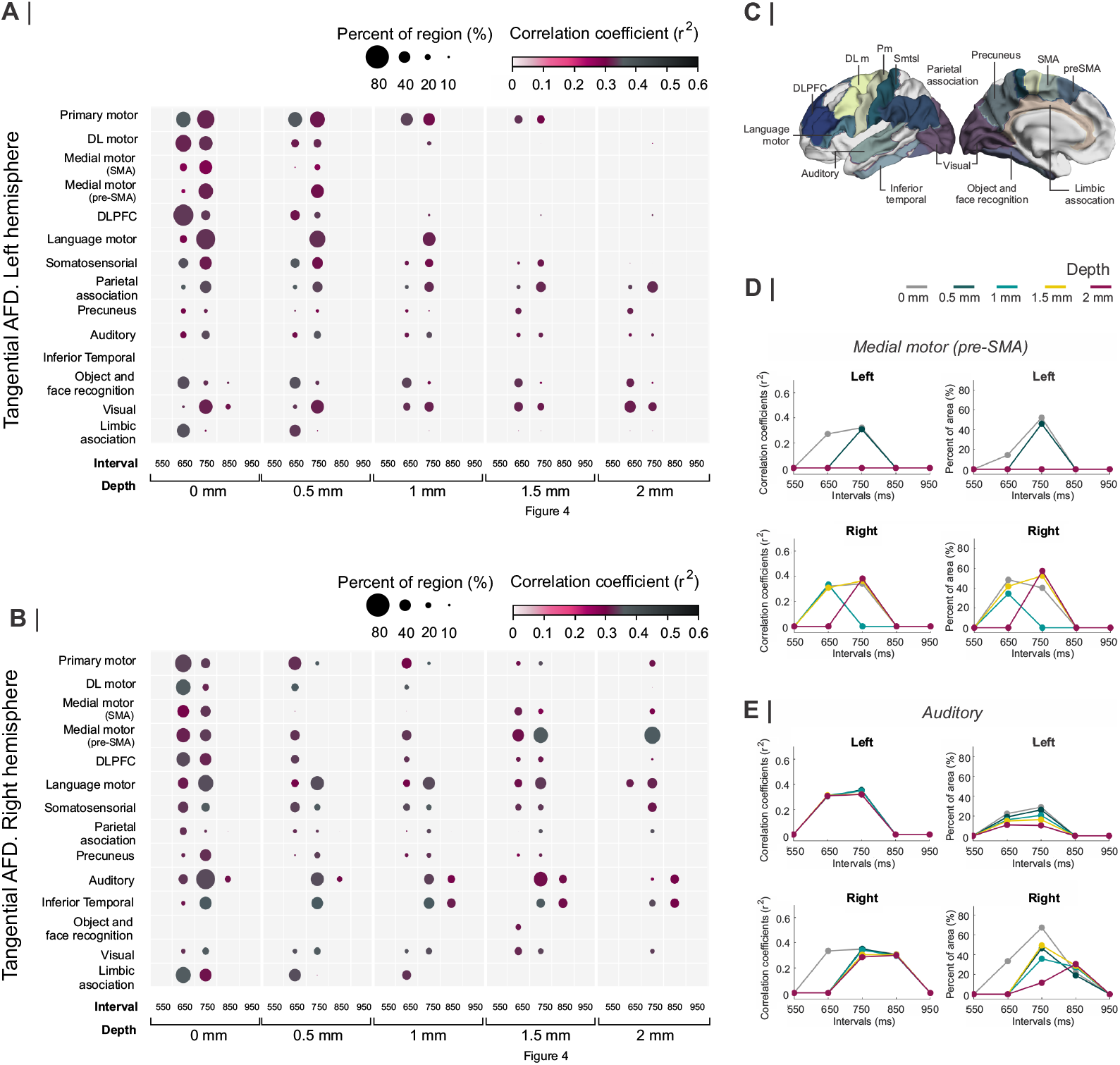
A and B. Interregional correlation plot of the auditory asynchronies and *_tan_AFD* as a function of the instructed interval and the depth of the *_tan_AFD* calculation. Significant clusters were aggregated into fourteen regions (y-axis) based on the Brainnetome atlas (C) (Fan et al., 2016). The colour and size of the circle for each cluster correspond to the correlation coefficient (critical value r^2^ > 0.355 at p < 0.02, df=29) and the percent of significant vertices in each area, respectively. The left hemisphere showed more areas with significant vertices than the right (17541 > 16641). A systematic decrease in circle size as a function of depth was observed across areas of both hemispheres. At a depth of 1.5 mm the regions with a larger percent of significant vertices for the right hemisphere at 750ms were the medial premotor (SMA and preSMA), auditory, and language motor areas (B), while small number of significant vertices were observed across the regions of the left hemisphere (A). In A and B it is possible to see the cluster analysis per area of the data from (Figure 4). D. Interval selectivity curves for the correlation coefficients and percent of significant vertices across *_tan_AFD* depths (colour coded) for preSMA. E: same and D but for auditory cortex. Note that the preferred interval in the two areas is between 650 and 750 ms.

To identify the anatomical regions with significant clusters of vertices, we parcellated the SWM based on the Brainnetome atlas (Fan et al., 2016). Areas with significative vertices (after correction for multiple comparisons: *p_FWE_* < 0.001 and *p_cluster_* < 0.001), were grouped into 14 regions: primary motor, dorsolateral secondary motor, medial motor (supplementary and pre-supplementary motor area), dorsolateral prefrontal cortex (DLPC), language motor, primary somatosensorial, parietal association, precuneus, primary and secondary auditory areas (auditory), inferior temporal cortex, object and face recognition areas, primary and secondary visual areas (visual), and limbic association areas (Figure 5C).

(Figure 5AB shows a systematic decrease in the percent of significant vertices as a function of depth across areas of both hemispheres. The most prominent effect is the interval selectivity of the correlation coefficients and percent of significant vertices, with a preferred interval between 650 and 750 ms, across many areas and depths (Figure 5ABDE). Specifically, at our target depth of 1.5 mm linked with the U-fiber system, the areas with the higher percent of significative vertices on the right hemisphere were pre-SMA (52.4%, r^2^=.41, p¡ 0.001), auditory (49.2%, r^2^=.38, p< 0.001) and language motor (40.2%, r^2^=.45, p< 0.001) for the correlation between the 750 ms auditory asynchronies and *_tan_AFD*. Similarly, at 650 ms the higher percent areas with significative vertices on the right hemisphere were the medial motor areas (pre-SMA, 41.8%, r^2^=.36, p< 0.001 and SMA 28.1%, r^2^=.39, p< 0.001) and language motor (28.1%, r^2^=.40, p< 0.001). Figures 5D and E show the interval selectivity curves for the correlation coefficients and percent of significant vertices across *_tan_AFD* depths (colour coded) for auditory cortex and preSMA, confirming the notion of interval preference for the behavioural and structural associations. In contrast, in the left hemisphere the percent areas with significant vertices were smaller, with significant effects for visual areas (31.8%, r^2^=.45, p< 0.001), object and face recognition (29.9%, r^2^=.44, p< 0.001) and primary motor areas (28.4%, r^2^=.44, p< 0.001) for the 650 ms interval; and parietal association (35.1%, r^2^=.52, p< 0.001) primary motor areas (28%, r^2^=.40, p< 0.001) and visual areas (25%, r^2^=.40, p< 0.001) for the 750 ms interval.

These results support the notion that the audiomotor system of the right hemisphere shows individual differences in the density of U-fibers with a gradient that correlates with the degree of prediction across subjects in the synchronization phase of the SCT. Crucially, this correlation is modality specific, suggesting that structural changes in the audiomotor system occur only for the modality that produces rhythmic timing with more precision, accuracy, and degree of predictability of the metronome (see Extended Data 2). Furthermore, the interval selectivity on the behavioural and structural correlation indicates that the timing circuits tuned at intervals of 650 and 750ms are the ones that are susceptible to present macro and microstructural differences in U-fibers of WM within the tested population of subjects. Finally, the visual areas and the motor cortex, contralateral (left hemisphere) to the tapping movements, also showed significant associations between rhythmic timing prediction and the U-fiber density.

### 2.4. Deep WM and structural chronotopic representation

We also evaluated the association between the precision, accuracy, prediction and error correction of rhythmic timing with the properties of the deep WM (dWM). Fixel-based analysis (FBA) (Dhollander et al., 2021; Raffelt et al., 2017) was used to estimate micro and macroestructural differences within-voxel of the dWM (Genc et al., 2018; Kelly et al., 2020; Mito et al., 2018; Rau et al., 2019). As with our analysis of superficial WM, this method is based on the constrained spherical deconvolution (CSD) of DWI data. FBA provides three fiber-specific indices (fiber density, fiber cross-section, and fiber density cross-section; FD, FC, and FDC, respectively) (Raffelt et al., 2017). FD is derived from the integral of the FOD lobes and is proportional to the total intra-axonal volume, thus reflecting the density of a population of fibers within a voxel (Rojas-Vite et al., 2019). If more than one fiber population coexist in a given voxel, the FOD is segmented, and a FD is assigned to each population referred to as fixel (fiber element). FC is a macroscopic metric of the fiber bundle diameter and, finally, FDC is a combination of FD and FC (see Methods). Briefly, the FBA analysis pipeline consists of five steps. First, the images of each subject are processed to obtain a WM Fiber Orientation Distribution (FOD) map in native space using CSD. Second, a template of the FODs and a fixel mask are built. Third, the AFD map and the corresponding fixels for each voxel are computed for each subject in native space. The FD metric is obtained for each fixel computed from the total DWI signal per voxel. Fourth the fixels and AFD map are reoriented to the template. Finally, fixel-wise statistics are performed at each spatial location in template space.

The FBA revealed significant negative correlations between the FDC in the corpus callosum (CC) and the predictive ability to synchronize to an auditory metronome: asynchronies for 650 ms, 750 ms, 850 ms and 950 ms intervals (Figure 6). Therefore, this analysis showed a tight relation between the density and bundle diameter of CC fibers and beat entrainment, with the subjects showing large predictive abilities and asynchronies close to zero also showing large FDC values, and the subjects with bad prediction and large asynchronies showing low FDC values (Extended Data 8). As in the case of the U-fiber metrics, the FBA values were not correlated with the other SCT behavioural parameters of the auditory and visual conditions during the synchronization and continuation epochs.

**Figure 6:**
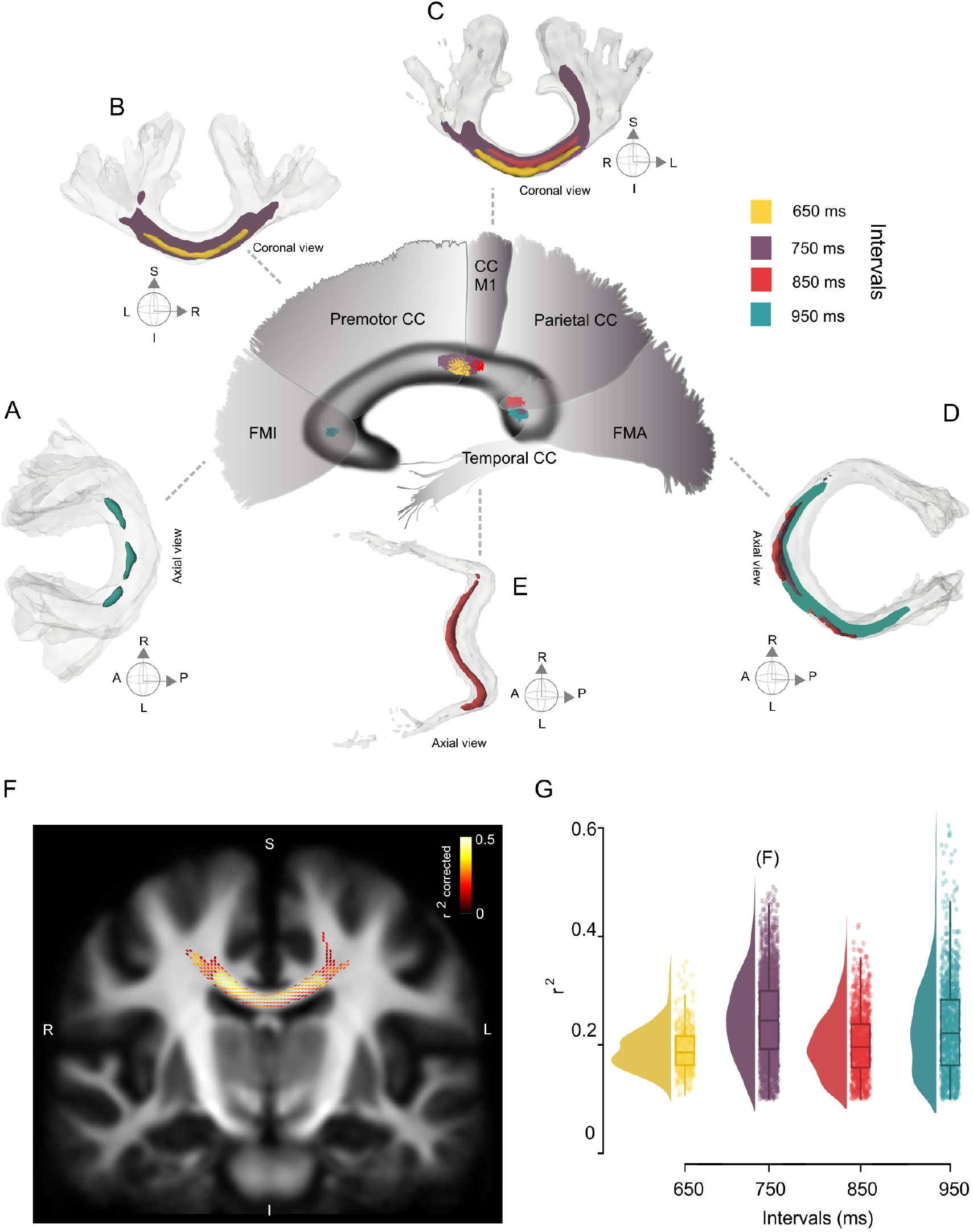
Chronotopic map of the correlation between the Deep White Matter (dWM). A-E. Fiber bundles that showed significant correlations between the asynchronies in the SCT auditory condition and the fiber density and cross-section (FDC) of the corpus callosum (CC). A chronotopic map with an anterior posterior gradient is evident. F. Coronal section of the CC showing the fixels with a significant correlation coefficient (colour coded r^2^ values; only fixels with pcorr<0.05 are shown) between asynchronies at the interval of 750 ms in the auditory condition and FDC. G. Distribution of correlation coefficient values (r^2^) of the FDC vs absolute asynchronies for the four intervals listed in the x-axis. On the right the interquartile box plots are depicted.

The association between rhythmic entrainment and white matter properties defined a chronotopic map in the CC, with FDC at different levels of CC showing significant correlations with the absolute asynchronies at specific intervals (Figure 6). This chronotopic map showed an anterior-posterior gradient, with behavioural and structural associations for short and long intervals in the anterior and posterior portions of the CC, respectively. Thus, the FDC fixel values of the posterior midbody of CC (interconnecting motor, pre-motor cortices and M1) showed a significant negative correlation with absolute auditory asynchronies for the 650 and 750 ms intervals (Figure 6BC; FWE-corrected P-value < 0.05). For the asynchronies at the intermediate interval of 850 ms, the negative correlation was observed with FDC fixel values located in the isthmus and the splenium (Figure 6C; interconnecting primary motor, temporal, and visual cortices). Finally, the asynchronies of the interval of 950 ms showed a significant negative correlation with fixels located in forceps minor (FMI) and major (FMA) (Figure 6A,D; interconnecting prefrontal and visual cortices, respectively).

It is evident in Figure 6A-E that the streamlines segments of the fixels with entrainment correlations were located mainly at the joint of the two hemispheres across the CC. Nevertheless, a lateralization effect was found for the left hemisphere, with fixels associated with auditory asynchronies of 750 and 950 ms in the isthmus and splenium, respectively. Besides the CC, right fornix showed a significant association with the asynchronies of the 750 ms interval for the FD metric (see Extended Data 8).

Lastly, we carried out a correlation analysis between the mean absolute asynchronies across the five intervals and the FC (Figure 7) and FDC (Extended Data 8). Notably, for the auditory condition the tracts with significant FC fixels were: left Arcuate Fasciculus (AF; Figure7A,E), CC M1 (Figure 7C,E), Forceps Mayor (FMA; Figure 7B,D), Superior Longitudinal Fasciculus 2 (SFL2; Figure 7F) and right Fornix (FX), (Extended Data 8).

**Figure 7:**
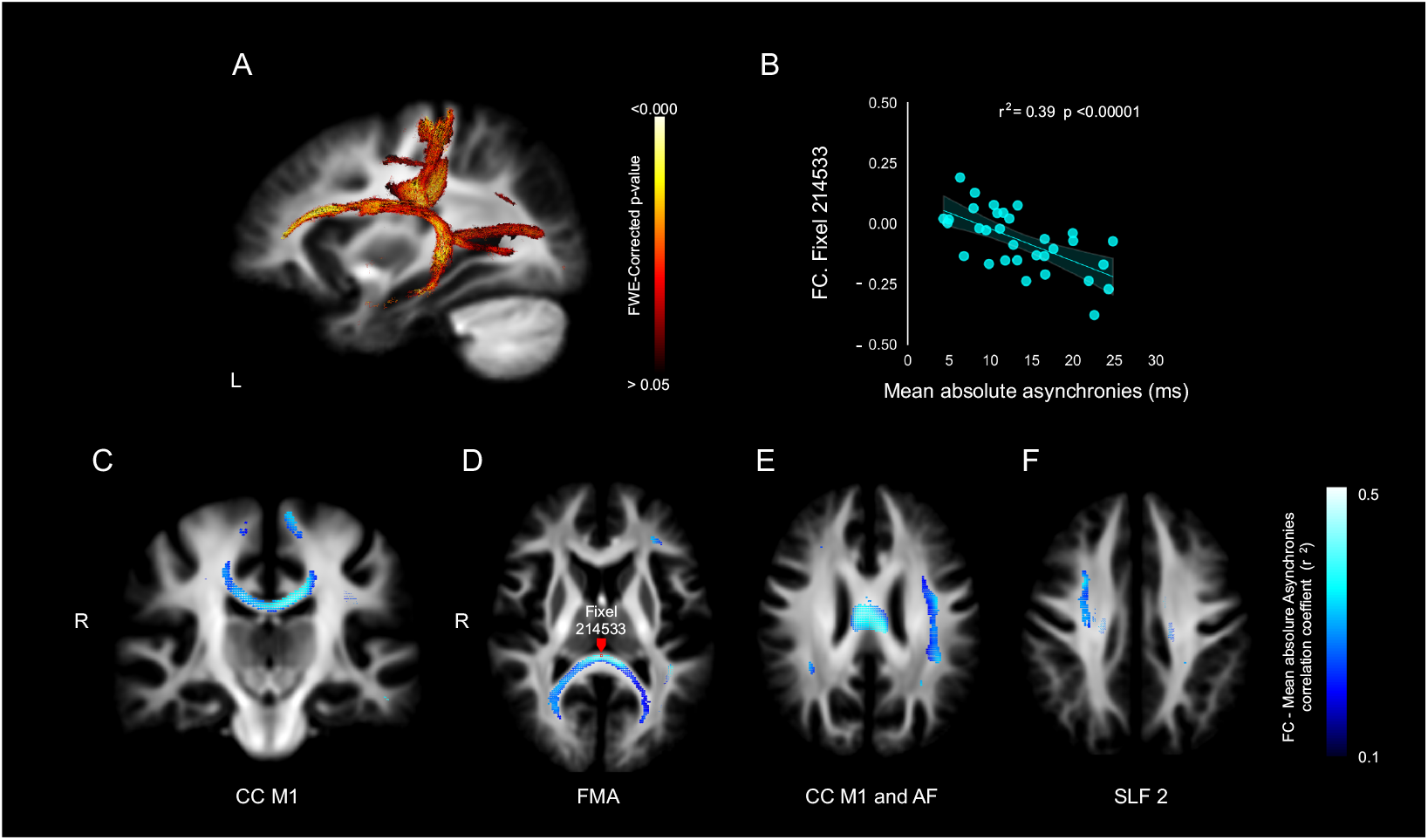
Association between Deep White Matter (dWM) and mean absolute asynchronies of the five tested intervals during SCT (550,650,750,850 and 950 ms). A. p-value of significant fixels after Family Wise Error (FEW) correction. B. Example of the significant correlation between the fiber cross-section (FC) metric and the mean absolute asynchronies in fixel 214533 (red dot in panel D). C, D, E and F. Negative correlation (r^2^) between FC and mean absolute asynchronies for significant fixels. CC= Corpus Callosum (C), FMA= Forceps Major (D), AF= Arcuate Fasciculus (E), SLF 2= Superior Longitudinal Fasciculus 2 (F).

## 3. Discussion

The present research determined the intersubject association between structural properties of the superficial and deep white matter and different measures of rhythmic timing during a synchronization-continuation tapping task. Our study supports five conclusions: First, healthy human subjects showed precision, accuracy, predictive entrainment, and error correction for rhythmic timing that was biased towards auditory instead of visual metronomes, confirming previous observations. Second, the right audiomotor system showed individual differences in the density of superficial white matter including the U-fibers that were correlated with the degree of predictive entrainment across subjects. Notably, these correlations were selective for the synchronization epoch of the auditory condition and were specific for the intervals of 650 and 750ms. Third, there was a significant association between predictive rhythmic entrainment and the density and bundle diameter of the corpus callosum, forming a chronotopic map with an anterior-posterior gradient. This implies that the behavioural and structural associations for short and long intervals were found in the anterior and posterior portions of the CC, respectively. Fourth, the fiber bundle diameter of the arcuate fasciculus, the Corpus Callosum, the Forceps Major and the Superior Longitudinal Fasciculus showed a significant correlation with the mean asynchronies across all tested tempos. Finally, no significant associations between the superficial and deep white matter properties were found with temporal variability, constant error, or lag one autocorrelation for the visual and auditory conditions during the synchronization and continuation epochs of the SCT. These last findings accentuate the fact that the micro and macro structural properties of the white matter are specifically linked with the predictive ability of the subjects to tap in synchrony with an auditory metronome.

Many studies have shown that performance tapping synchronized to an auditory metronome is more precise and accurate than synchronization to a flashing visual metronome with the same timing characteristics (Chen et al., 2002; Hove et al., 2013; Merchant et al., 2008c; Patel et al., 2005; Repp & Penel, 2004; Zarco et al., 2009).Since the first processing relays, the auditory system has higher temporal resolution compared to the visual system (Duysens J. & A., 1996; He et al., 1997; Sayegh et al., 2011), and plays a critical role on time perception and reproduction across many tasks, not only on tapping SCT (Grondin et al., 2005; Merchant et al., 2008b; Merchant & de Lafuente, 2014). For example, when audiovisual stimuli are used in an oddball paradigm, the perceived duration is dominated by the auditory modality (Chen & Yeh, 2009). In addition, TMS disruption of the auditory cortex impaired not only time estimation of auditory but also of visual stimuli, while disruption in the visual cortex only produced timing impairments for visual stimuli (Kanai et al., 2011). Consequently, a current hypothesis is that the auditory cortex is engaged in multimodal temporal processing and the interaction between the auditory and the motor system in the frontal lobe allows not only for time encoding but also time prediction (Merchant & Honing, 2014; Merchant & Yarrow, 2016; Patel & Iversen, 2014; Schwartze et al., 2012). Our present findings support this notion in different ways. First, the rhythmic timing performance of our subjects was biased towards the auditory condition across different behavioural measures. Second, the right audiomotor system showed widespread and significant correlations between the density of superficial U-fibers and the degree of auditory predictive entrainment across subjects. These anatomo-behavioural associations are selective to the intervals in the 650 to 750 range. Third, the fiber bundle diameter of the left arcuate fasciculus, a key tract connecting the parieto temporal auditory system with the frontal lobe, showed a significant correlation with the mean asynchronies across all tested tempos. The later results are in line with a recent study that showed that the left arcuate fasciculus show altered connectivity and fractional anisotropy reduction in stuttering subjects compared with fluent speakers (Jossinger et al., 2022). Furthermore, the clear clustering of subjects as either good or bad synchronizers in a syllable isochronous entrainment task correlates with both the difference in the activation magnitude in frontal areas and changes in the white matter pathways (i.e. left arcuate fasciculus) that connect the auditory with the premotor cortical system (Assaneo et al., 2019). Hence, our results accentuate the audiomotor structural foundation for rhythmic entrainment (Honing & Merchant, 2014; Miyata et al., 2022). The lack of structural associations in the visual condition is probably due to the larger variability in rhythmic tapping for this modality, making difficult to infer statistical correlations between our metrics.

The observed associations between the measured properties of superficial and deep white matter and the rhythmic entrainment prediction were negative, indicating that the subjects with asynchronies close to zero and large predictive abilities showed large values in the white matter parameter, while the subjects with bad prediction and large asynchronies showed low values on the same metrics. These results support the theory that intersubject differences in rhythmic entrainment prediction depend on micro and macrostructural white matter properties, which could have a genetic and/or learned substrate. On the genetic side, we could speculate that the existence of poor and superior synchronizers (Blecher et al., 2016) may depend on the fiber density of superficial U-fibers in the right audiomotor system as well as on the density of deep tracts such as the CC and the arcuate fasciculus. On the training side, these superficial and deep white matter bundles may develop larger density and myelination during intense musical practice, distinguishing the audiomotor tracts between musicians and non-musicians (Palomar-García et al., 2020; Vaquero et al., 2018; Zatorre et al., 2007).

The correlations between the density of tangential U-fibers in the right audiomotor circuit and the asynchronies for the auditory condition were interval selective for the intermediate tested tempos. This interval specificity supports the idea that the spontaneous rhythmic tempo observed in many studies of rhythmic entrainment, and with values between 600 and 750 ms, (Drake et al., 2000a,b; McAuley et al., 2006) (see (Figure 2D) has its biological origins in the structural properties of the U-fibers running superficially across the audiomotor circuit. In this scenario, the cortico-cortical connectivity within the audiomotor system is specially designed to support the internal representation of pulse at the preferred tempo, providing larger predictive abilities across subjects for the spontaneous motor tempi (Balasubramaniam et al., 2021). Further anatomofunctional studies are required to test this hypothesis.

Ultra-high field (7 T) functional imaging revealed that the medial premotor areas (SMA-preSMA) of the human brain possess neural circuits that are tuned to different durations forming a topographical arrangement. This chronomaps show units with enhanced responses for the preferred interval and suppressed activity for the nonpreferred duration, and define a topographical gradient with short preferred intervals in the anterior portion and long preferred intervals in the posterior portion of the medial premotor areas (Protopapa et al., 2019; Schwartze et al., 2012). Our measurements on the FDA also revealed a topographical arrangement in the correlations between the density of deep fibers and in CC and the rhythmic predictive abilities of subjects, so that significant anatomobehavioural associations were found in the anterior part of the CC for short intervals and in the posterior CC for long tapping tempos (Schwartze et al., 2012). Nevertheless, our chronotopic map is defined by the correlation between the asynchronies and the FDA, with no functional basis. Moreover, it extends from the CC anterior portions that connect the two hemispheres of the premotor areas of the frontal cortex all the way to the occipital lobe. Hence, the present topography goes well beyond SMA. In line with our present results, a recent imaging study described the existence of large chronomaps covering the cortical mantle from the dorsal and ventral premotor areas to the occipital pole (Harvey et al., 2020; Hendrikx et al., 2022). Consequently, timing maps define a cortical processing framework for efficient timing integration that has both functional and anatomical basis.

Previous studies have shown correlations between the performance of SMS tasks and microstructural characteristics of deep white matter (DWM) derived from diffusion-weighted magnetic resonance images (DWI). Blecher (Blecher et al., 2016) found significant correlations between asynchronies of participants that performed a finger-tapping task with an auditory metronome in the millisecond scale and fractional anisotropy (FA) of the left arcuate fasciculus and in the temporal segment of the corpus callosum. Better performance in the finger-tapping task was associated with higher values in FA. Nevertheless, the superficial white matter (SWM) immediately below the cortex has so far been neglected, despite its volume (60 % of the total WM volume) and its proximity to neocortical regions having a pivotal role in maintaining cortico-cortical connectivity (Schüz & Braitenberg, 2002; Schmahmann et al., 2006). The SWM is integrated for short-range association fibers that connect adjacent gyri (U-fibers) as well as for the initial or final part of the projection and association (large-range) tracts that form the deep white matter(Schüz & Braitenberg, 2002; Guevara et al., 2020; Kirilina et al., 2020; Yoshino et al., 2020). In recent years abnormalities of SWM have been described in epilepsy (Liu et al., 2016), autism spectrum disorder (Hong et al., 2019), Alzheimer ’s disease (Phillips et al., 2016), schizophrenia (Nazeri et al., 2013), and stroke (Stockert et al., 2021). Two limitations of studying the SWM in vivo have been constantly described: the fiber-crossing and lower values of anisotropy in comparison with the dWM (Guevara et al., 2020). In order to solve that, we joined two methods for the study of the SWM using synthetic streamlines to query the properties of SWM at different depths and the FD values associated with different orientations of the fiber. This approximation allowed us to perform sub-voxel analyses at the fiber-specific level, describing local macrostructure and microstructural properties. FBA, on the other hand, has been used to evaluate deep white matter in the context of healthy aging, childhood development, psychiatric disorders, fetal and neonatal development, and other disorders (Dhollander et al., 2021). Our results showed that the subjects’ accuracy in the performance of SCT was associated with higher FC, FDC or FD values. This is in accordance with the literature that shows that better performance in different tasks is associated with higher values of FBA metrics. For example, better performance in a task of bimanual coordination was associated with higher values of the FBA in the corpus callosum (Adab et al., 2020). Furthermore, a global decrease in FBA has been observed in studies on the aging effect (Choy et al., 2020), memory decline in older and younger adults (Radhakrishnan et al., 2020), neurodegenerative and demyelinating disorders (Al-Amin et al., 2020; Carandini et al., 2021).

## Acknowledgements

We thank Sonja Kotz and Florencia Assaneo for their fruitful comments on the manuscript. We also thank Luis Prado, Erick Pasaye, Juan Ortiz, Leopoldo González, for their technical assistance.

## Methods

### Participants

Thirty-two healthy human subjects (age= 25.37 ± 3.21 years; 19 females) without musical training volunteered to participate and gave informed consent, which complied with the Declaration of Helsinki and was approved by our Institutional Review Board. All participants were right-handed and native Spanish speakers. They did not have MRI contraindications nor neurological, psychiatric, or cognitive conditions.

### Apparatus

Participants were seated comfortably in a quiet experimental room, facing a high-definition 23” touch screen (refresh rate: 60 Hz; ELO Touch solutions) located 50 cm away, which they were instructed to tap using the right index finger. Auditory stimuli were presented through noise-cancelling headphones (Sony, MDR-NC50).

### Experimental Task

The SCT of the present study is similar to the standard SCT described elsewhere(Merchant et al., 2008c,b) however, instead of tapping a button, the subject tapped on the right or left halves of the touch screen. The task started when three empty white circles (radius 1 degree of visual angle) on a black background were presented simultaneously, forming an inverted isosceles triangle (2 degrees of visual angle on each side). Subjects were trained to place their finger at central-bottom target to start the trial, and to attend to a sequence of six right/left alternating stimuli with a constant interstimulus interval (perception epoch), tapping the touch screen on the corresponding target in synchrony with metronome during six intervals (synchronization epoch), and to continue taping on the screen for other six intervals without the metronome (continuation epoch) (Figure 1). The mean asynchronies were presented at the end of the trial as feedback. A trial was considered correct when the asynchronies were below 30% of the target interval during the synchronization and below 40% during the continuation epoch. The task was programmed using Psychtoolbox for Matlab (2018, The Mathworks).

### Stimuli

The visual metronome (33 ms, yellow circle, 0.5 degrees of visual angle) was alternatively displayed inside of the right or left empty circles. The auditory metronome (33 ms, 550 Hz, 80–85 dB SPL), was also presented alternatively on the right or left headphone. The isochronous target intervals were 550, 650, 750, 850, and 950 ms, which were pseudorandomly presented within a block. Each subject performed 10 correct trials for each target interval.

### Procedure

The MRIs were collected in the first session. In a second visit, outside of the MRI scanner, the auditory and visual tasks were performed randomly across subjects.

### Analysis of behavioral data

Four metrics were calculated to assess the subjects’ performance during SCT (Figure 2A) (Gámez et al., 2018; Merchant et al., 2008c). During the synchronization epoch, we estimated the absolute asynchronies and autocorrelation of the inter-tap interval time series (Iversen et al., 2015; Wing, 2002). The constant error and temporal variability were calculated from the produced intervals during both the synchronization and continuation epochs. Absolute asynchronies were defined as the unsigned difference between stimulus onset and the tap onset. Constant error was calculated as the difference between the produced and the target interval and is a measure of timing accuracy. Temporal variability was defined as the standard deviation of the total produced intervals and is a metric of timing precision. The autocorrelation of the six inter-tap-intervals during a trial was calculated and averaged across trials. Thus, lag 1 autocorrelation is normally negative for isochrone metronomes, meaning that a longer produced interval tends to be followed by a shorter interval and vice versa, reflecting an error correction mechanism used to not lose the beat of the metronome during synchronization (Iversen et al., 2015). A repeated-measures ANOVA, with two and three factors, was carried out for the analysis of asynchronies, constant error and temporal variability.

### Imaging protocol

Images were acquired using a 3 T Philips Achieva TX scanner with a 32- channel head coil. T1-weighted volumes were obtained using a three-dimensionally encoded spoiled gradient echo sequence (TR/TE=8.2/3.7 ms, flip angle = 8°, field of view = 256×256×176 mm3, matrix size = 256×240×176 mm3 yielding voxel resolution = 1×1×1 mm3). Diffusion-weighted images (DWI). Diffusion-weighted images (DWI) were acquired with echo-planar imaging (EPI), 2×2×2 mm3 voxel resolution, FOV = 256 × 256 mm3, 62 axial slices, TR/TE=16500/72 ms. Images were sensitized to diffusion with b=1000 s/mm^2^ (64 unique directions) and b=3000 s/mm^2^ (96 directions). Five volumes without diffusion weighting (b = 0 s/mm2) were also acquired, along with an additional b = 0 s/mm2 volume obtained with reversed-phase encoding polarity for correction of geometric distortions.

### Image processing

a. Cortical surface T1-weighted images were used to estimate cortical surfaces. Images were first denoised (Coupe et al., 2008), and corrected for intensity inhomogeneities (Tustison et al., 2011). Binary masks of the brain were obtained with volBRAIN v.2.0 (Manjón & Coupé, 2016). Cortical surfaces were obtained through the FreeSurfer pipeline v.6.0 (Fischl, 2012). Individual surfaces were registered to the surface template with 20,484 vertex (fsaverage5).
b. Diffusion-weighted volumes (DWI) were first denoised (Manjon et al., 2013) and bias field corrected (Tustison et al., 2011), then corrected for geometric distortions and motion using FSL ’s topup-eddy algorithm (Andersson & Sotiropoulos, 2016).

### Fixel-based analysis (FBA)

We analyzed individual fiber-specific properties in the presence of crossing fiber populations (‘fixels’) (Raffelt et al., 2015), following the steps described in (Raffelt et al., 2017), using the tools available in MRtrix3 (Tournier et al., 2019). A WM mask was computed for each subject, followed by global signal intensity normalization of the DWI, which was performed across subjects by dividing all volumes by the median b = 0 s/mm2 intensity. Images were upsampled to 1 mm3 isometric resolution. WM Fiber Orientation Distributions (FODs) were estimated using the Multi-Shell Multi-Tissue Constrained Spherical Deconvolution (MSMT-CSD) algorithm (Jeurissen et al., 2014). Tissue-specific response functions were calculated for each subject, from which we derived group-averaged response functions that were used to estimate FODs (lmax=8) for each subject. A template of FODs was constructed through iterative nonlinear registration, using the FODs of all 32 subjects followed by the calculation of the intersection of masks of all subjects. Fixels were derived at each voxel by segmentation of the FODs and reoriented to the study template. Finally, FBA metrics (fiber density, fiber cross-section, and fiber density cross-section; FD, FC, and FDC, respectively) were calculated for each fixel. FBA Statistical analysis. A whole-brain probabilistic tractogram was calculated based on the FODs population template, seeded from a whole-brain white matter mask to produce a tractogram of 20 million streamlines. Next, the SIFT algorithm (Smith et al., 2013) was used to select a subset of streamlines (n=2 million) that best fit the diffusion signal and therefore reduce tractography biases. Based on the probabilistic tractography, the structural connectivity metric between fixels was obtained by performing the Connectivity-based fixel enhancement (CFE) tool (Raffelt et al., 2015).

The measures of FD, FC, and FDC were correlated with the performance in the SCT for both conditions (visual and auditory) using a General Linear Model. Family-wise error (FWE) permutation test was carried out for multiple comparisons correction (Nichols & Holmes, 2001).

Following statistical analysis, the identification of the tracts with significant fixels in the group space was done using the tract-selection regions from XTRACT tool (Warrington et al., 2020) included in FSL software (FMRIB’s Software Library – FSL, Oxford, U.K) (Smith et al., 2004), which were warped into our population template. Tracts with significant fixels were reconstructed using MRTrix (Tournier et al., 2019).

### Superficial White Matter surfaces

To assess the properties of SWM we used a synthetic representation of axons based on cortical topology. We computed a Laplacian potential field between the ventricles and the GM-WM boundary (Jones et al., 2000; Lerch et al., 2008; Liu et al., 2016) using minclaplace (Lerch et al., 2008). Next, Laplacian-streamlines were seeded at each vertex of the WM surface and propagated towards the ventricles using Matlab, 2020A. Thus, one Laplacian streamline was obtained for every WM surface vertex (Figure 3A). The distance between each step of the Laplacian-streamline was 100 μm, and streamlines were truncated at 5 mm. This resulted in smooth and non-overlapping pathways that respect topology and traverse the superficial white matter. Furthermore, the first segment of these streamlines is perpendicular to the GM-WM surface, with subsequent segments gradually bending as they extend away from it. These aspects make the Laplacian streamlines behave similarly to what is expected from the anatomy of the SWM. Finally, all the WM surfaces and synthetic streamlines were warped to their corresponding subject-specific DWI space via between-modality non-linear registration using ANTS (Avants et al., 2011).

We used the fixel-based information to independently evaluate the two fiber systems that coexist in the SWM: U-fibers subserving short-range cortico-cortical connectivity, and long-range projection, association and commisural fibers (Kirilina et al., 2020; Schüz & Braitenberg, 2002; Yoshino et al., 2020). With the assumption that U-fibers run tangentially to the GM-WM surface, while long-range fibers impinge on the surface in a perpendicular fashion, we attributed the fixels oriented parallel to the segments of the Laplacian streamlines to long-range fibers, and the remaining fixels to the U-fiber system. DWI-derived metrics were sampled along each Laplacian streamline at 0,0.5,1,1.5 and 2 mm under the GM-WM interface. The metrics sampled were *_total_AFD* (the integral of the FODs within a given voxel), and AFD attributed to either long-range fibers (*_par_AFD*; the integral of the FOD of the fixel parallel to the Laplacian streamline), or U-fibers (*_tan_AFD*; defined as *_total_AFD* - *_par_AFD*).

All metrics along the Laplacian streamlines were projected onto the GM-WM surface of the fsaverage5 template for visualization and statistical analyses and smoothed using a two-dimensional kernel of 15 mm of full width at half maximum.

### Surface-based analysis of SWM

Analyses were performed by fitting a general linear model at each vertex. This analysis assessed the relation between the value of diffusion metrics in each vertex (_*i*_) and the behavioral metrics from the SCT (absolute asynchronies, constant error, temporal variability, and lag 1 of the autocorrelation of the inter-tap-interval time series), as:

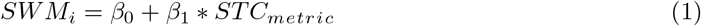

Surface vertex-wise analysis was corrected for multiple comparitions using random-field theory with a threshold of family-wise error to *(p_FWE_* < 0.001). Clusters with *p_cluster_* < 0.001 were deemed significant.

Significant clusters were anatomically identified using on the Brain Atlas Based on Connectional Architecture (Brainnetome) (Fan et al., 2016). All the analyses were carried out in Surfstat (Worsley et al., 2009) for Matlab (2018; The Mathworks).

## 4. Extended data

**Extended data 1:**
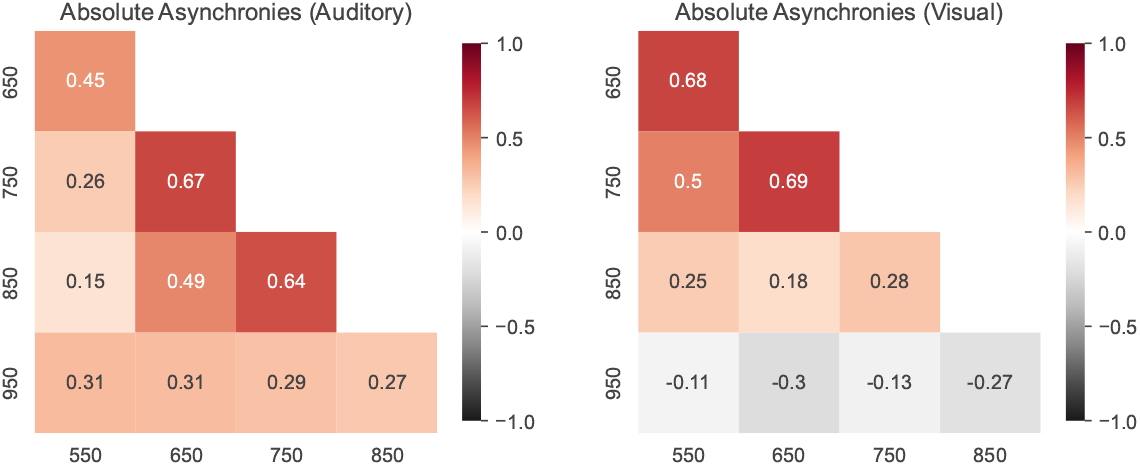
Correlation matrix for the Intersubject absolute asynchronies across instructed intervals for the auditory (left) and visual (right) conditions. Note that only the correlation between the intervals of 650 and 750 ms was significant for both the auditory and visual conditions.

**Extended data 2:**
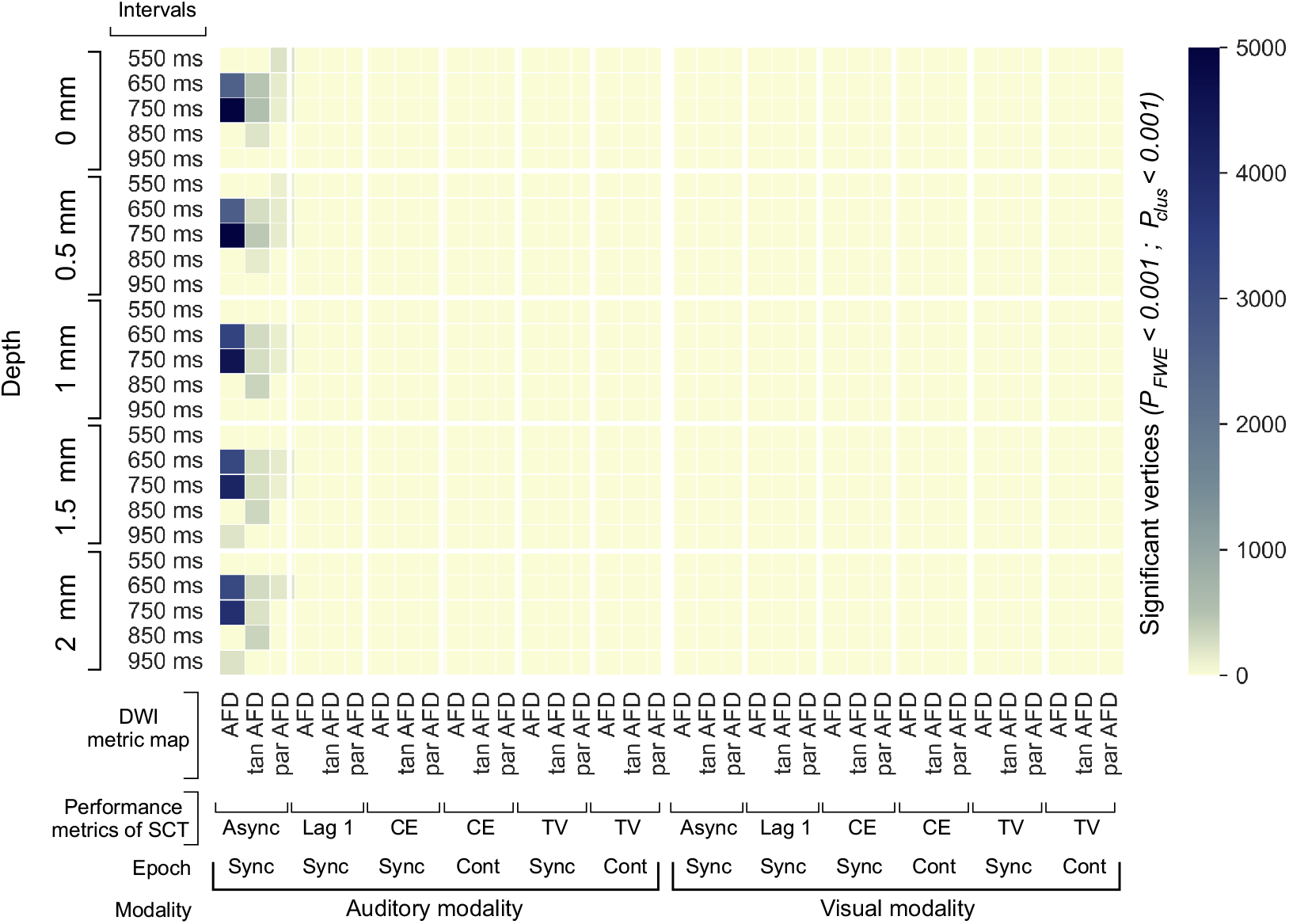
Characterization of the SWM properties associated with the performance of SCT. The Surface-based analysis (SBA) was used to assess the association between the performance metrics of the SCT and the metrics to characterize the SWM sampled in five depths with respect to the grey/white matter interface. The number of significant vertices (both hemispheres) after SBA are shown for the three DWI metric maps, through five depths sampled and the four behavioral metrics, in five intervals, two epochs, and two modalities. The number of vertices for the three AFD metrics showed association only with the absolute asynchronies of the different intervals of the auditory modality. The largest significant clusters were observed between the absolute asynchronies for the 650 and 750 ms intervals through the five depths sampled in the three AFD metrics. The SBA results were corrected to multiple comparisons, using as thresholds *P_FWE_* < 0.001 and *P_cluster_* < 0.001. AFD (Apparent Fiber Density), *_tan_AFD* (Tangential Apparent Fiber Density), *_par_AFD* (Parallel Apparent Fiber Density). Asynch (absolute asynchronies), Lag 1 (Lag 1 of the autocorrelation for each instructed interval), CE (Constant Error), TV (Temporal Variability), Sync (Synchronization), Cont (Continuation).

Extended data 3. Video

Extended data 3. Spatial correlations between the SWM Apparent Fiber Density and interval-specific absolute asynchronies of the auditory modality. Correlation coefficients (r^2^) of the significant cluster vertices after surface-based analysis through the depths sampled (p_*FWE*_ < 0.001 and p_*cluster*_ < 0.001).

Extended data 4. Video

Extended data 4. Spatial correlations between the Tangential SWM Apparent Fiber Density and interval-specific absolute asynchronies of the auditory modality. Correlation coefficients (r^2^) of the significant cluster vertices after surface-based analysis through the depths sampled (p_*FWE*_ < 0.001 and p_*cluster*_ < 0.001).

Extended data 5. Video

Extended data 5. Spatial correlations between the Parallel SWM Apparent Fiber Density and interval-specific absolute asynchronies of the auditory modality. Correlation coefficients (r^2^) of the significant cluster vertices after surface-based analysis through the depths sampled (p_*FWE*_ < 0.001 and p_*cluster*_ < 0.001).

**Extended data 6:**
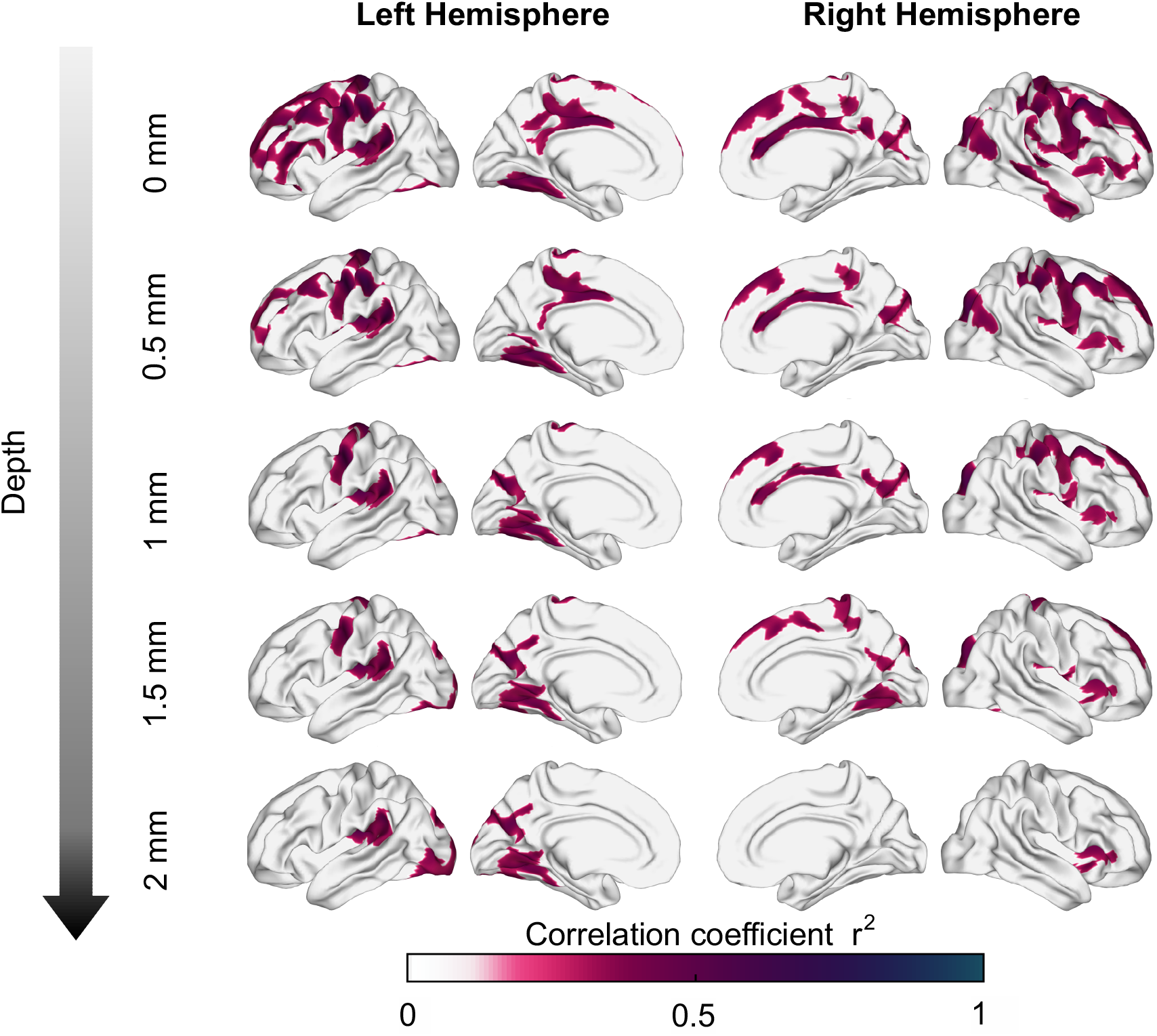
Association between asynchronies (650 ms intervals) and tangential superficial white matter apparent fiber density at various depths from the GM/WM interface. *_tan_AFD* sampled at 0 mm shows significant correlations distributed throughout most of the brain, but specific regions (e.g., motor, auditory and visual areas) consistently show significant correlations as depth increases.

**Extended data 7:**
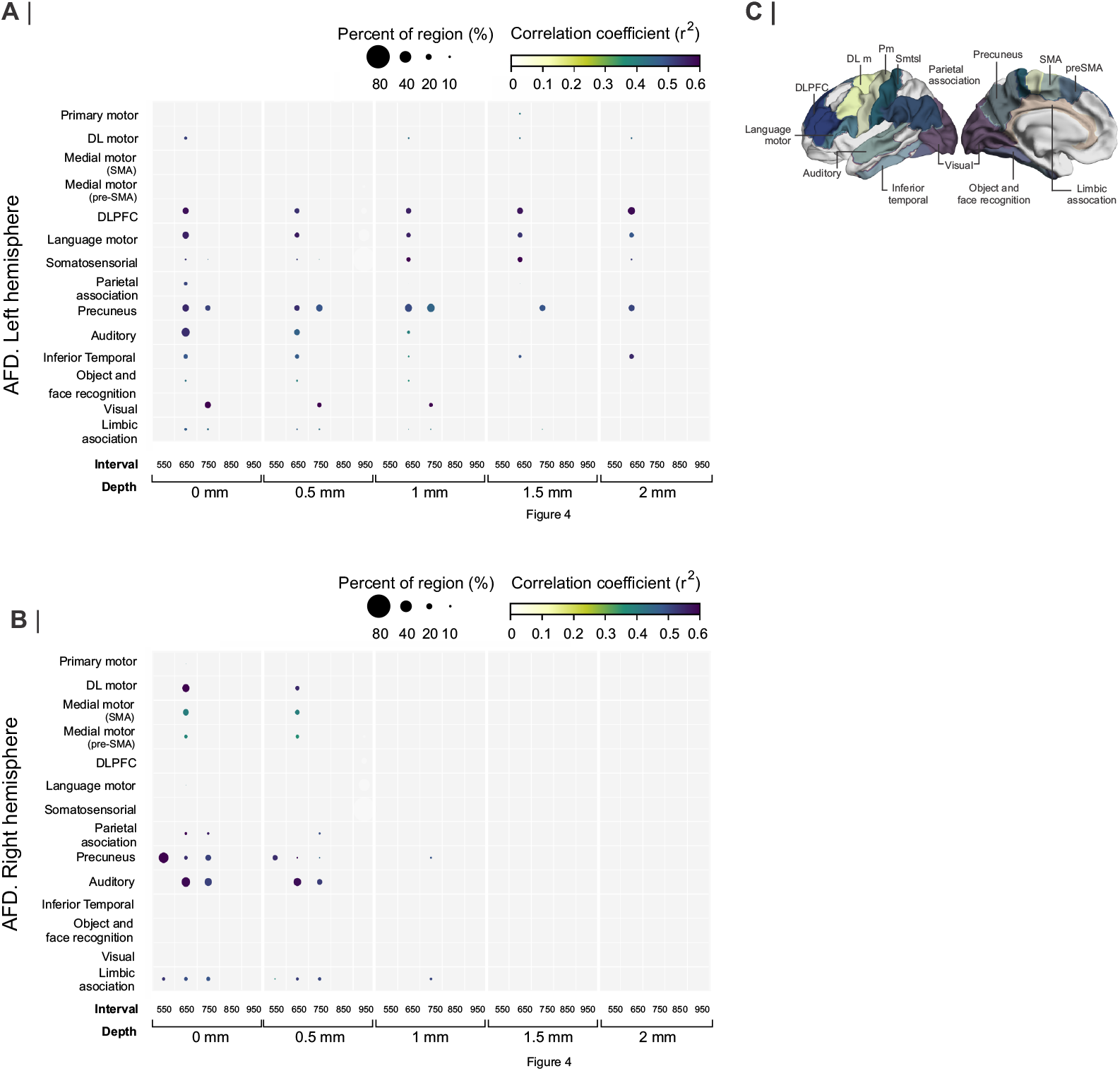
A and B. Interregional correlation plot of the auditory asynchronies and *_par_AFD* as a function of the instructed interval and the depth of the *_par_AFD* calculation.

**Extended data 8:**
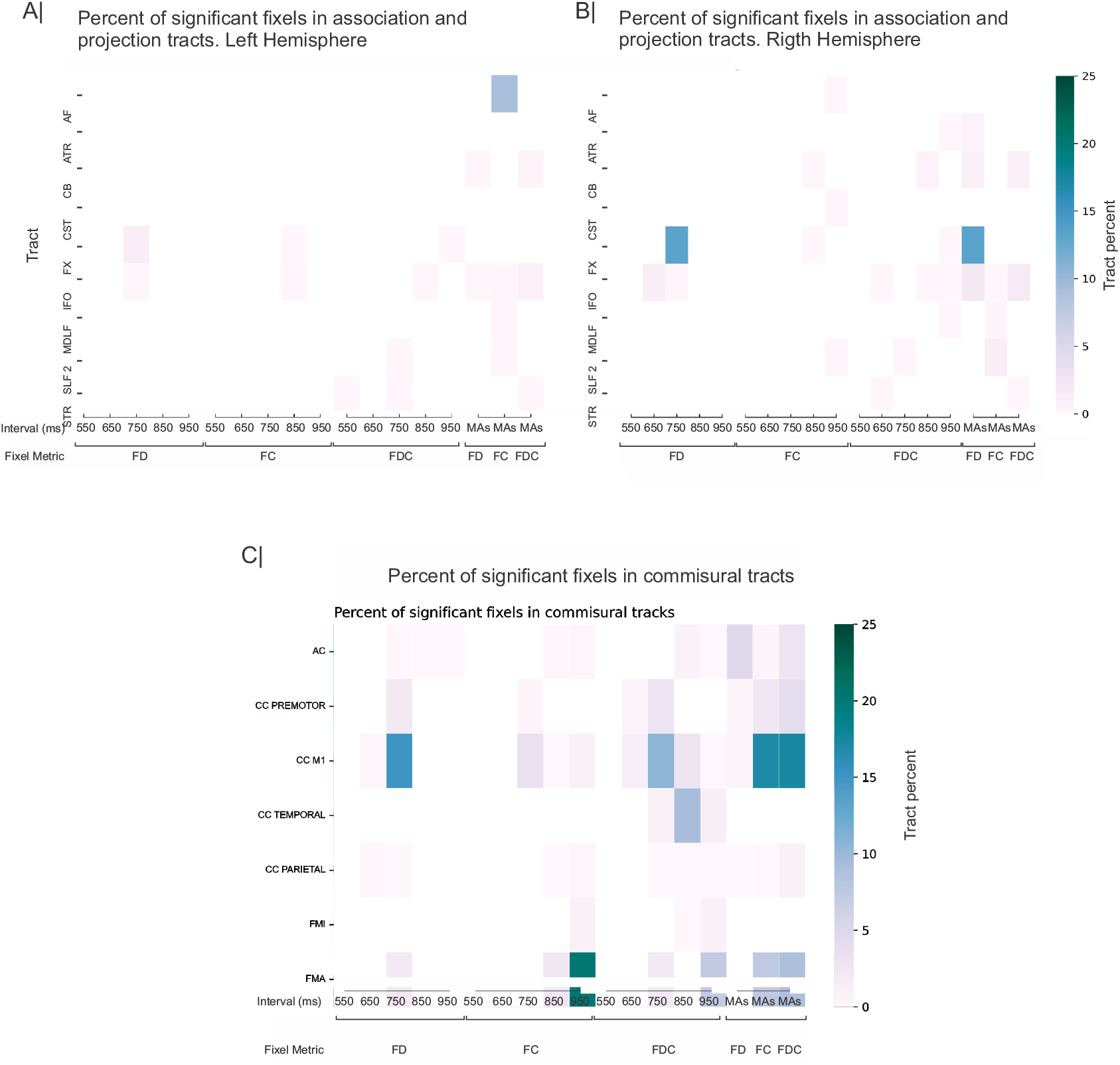
Percent of significative fixels in the Deep White Matter in projection tracts (A) Left, (B) Right and in commissural tracts (C). AF= Arcuate fasciculus; ATR = Anterior Thalamic Radiation ; CB = cingulum bundle; CST= Corticospinal Tract ; FX = fornix; IFO= Inferior Longitudinal; MDLF= Middle Longitudinal Fascicle ; SLF 2= Superior Longitudinal Fasciculus 2; STR ; AC= Anterior commisure; CC = Corpus Callosum; FMI= Forceps Minior; FMA= Forceps Major; FD= Fiber Density; FC = Fiber Cross; FDC= Fiber Density and Cross; MAs = Mean Asychronies.

