## Extended data for "White matter structural bases for predictive tapping synchronization"

### 1. Extended data

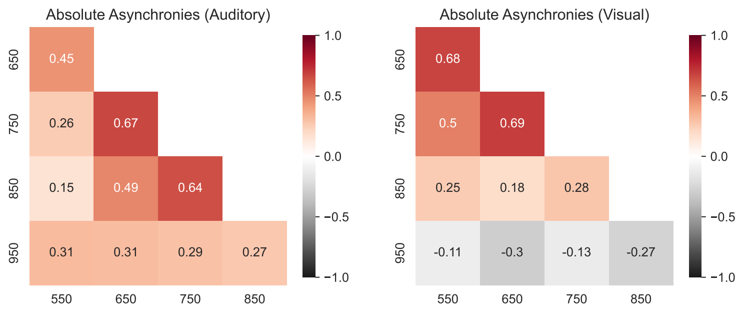


Extended data 1: Correlation matrix for the Intersubject absolute asynchronies across instructed intervals for the auditory (left) and visual (right) conditions. Note that only the correlation between the intervals of 650 and 750 ms was significant for both the auditory and visual conditions.

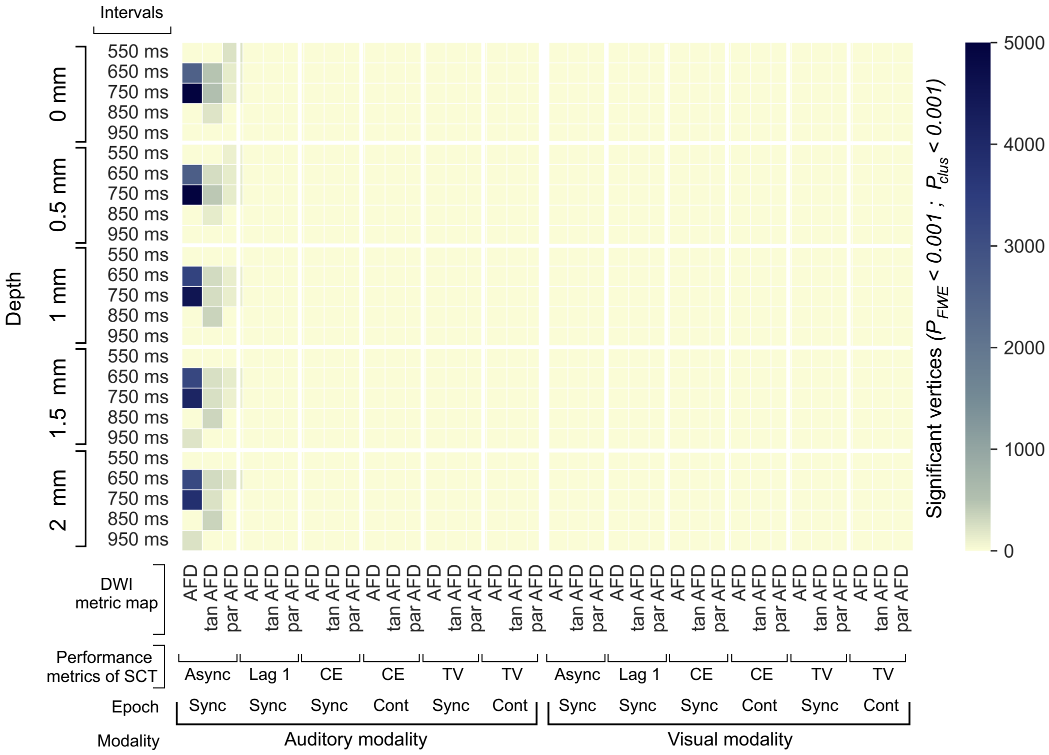


Extended data 2: Characterization of the SWM properties associated with the performance of SCT. The Surface-based analysis (SBA) was used to assess the association between the performance metrics of the SCT and the metrics to characterize the SWM sampled in five depths with respect to the grey/white matter interface. The number of significant vertices (both hemispheres) after SBA are shown for the three DWI metric maps, through five depths sampled and the four behavioral metrics, in five intervals, two epochs, and two modalities. The number of vertices for the three AFD metrics showed association only with the absolute asynchronies of the different intervals of the auditory modality. The largest significant clusters were observed between the absolute asynchronies for the 650 and 750 ms intervals through the five depths sampled in the three AFD metrics. The SBA results were corrected to multiple comparisons, using as thresholds $P_{FWE} < 0.001$ and $P_{cluster} < 0.001$. AFD (Apparent Fiber Density), $_{tan}AFD$ (Tangential Apparent Fiber Density), $_{par}AFD$ (Parallel Apparent Fiber Density). Asynch (absolute asynchronies), Lag 1 (Lag 1 of the autocorrelation for each instructed interval), CE (Constant Error), TV (Temporal Variability), Sync (Synchronization), Cont (Continuation).

Extended data 3. Video

Extended data 3. Spatial correlations between the SWM Apparent Fiber Density and interval-specific absolute asynchronies of the auditory modality. Correlation coefficients ($r^2$) of the significant cluster vertices after surface-based analysis through the depths sampled ($p_{FWE} < 0.001$ and $p_{cluster} < 0.001$).

Extended data 4. Video

Extended data 4. Spatial correlations between the Tangential SWM Apparent Fiber Density and interval-specific absolute asynchronies of the auditory modality. Correlation coefficients ($r^2$) of the significant cluster vertices after surface-based analysis through the depths sampled ($p_{FWE} < 0.001$ and $p_{cluster} < 0.001$).

Extended data 5. Video

Extended data 5. Spatial correlations between the Parallel SWM Apparent Fiber Density and interval-specific absolute asynchronies of the auditory modality. Correlation coefficients ($r^2$) of the significant cluster vertices after surface-based analysis through the depths sampled ($p_{FWE} < 0.001$ and $p_{cluster} < 0.001$).

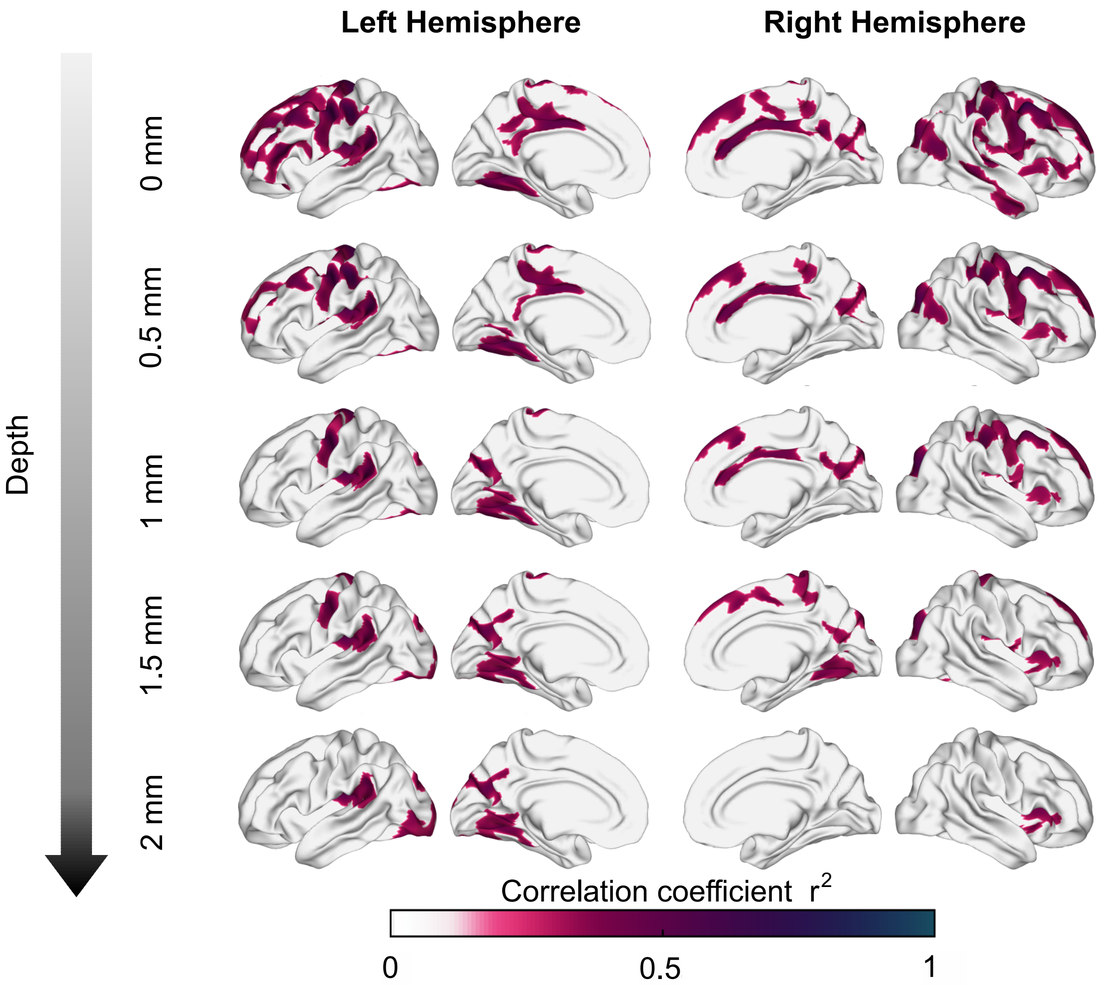


Extended data 6: Association between asynchronies (650 ms intervals) and tangential superficial white matter apparent fiber density at various depths from the GM/WM interface. ${}_{tan}AFD$ sampled at 0 mm shows significant correlations distributed throughout most of the brain, but specific regions (e.g., motor, auditory and visual areas) consistently show significant correlations as depth increases.

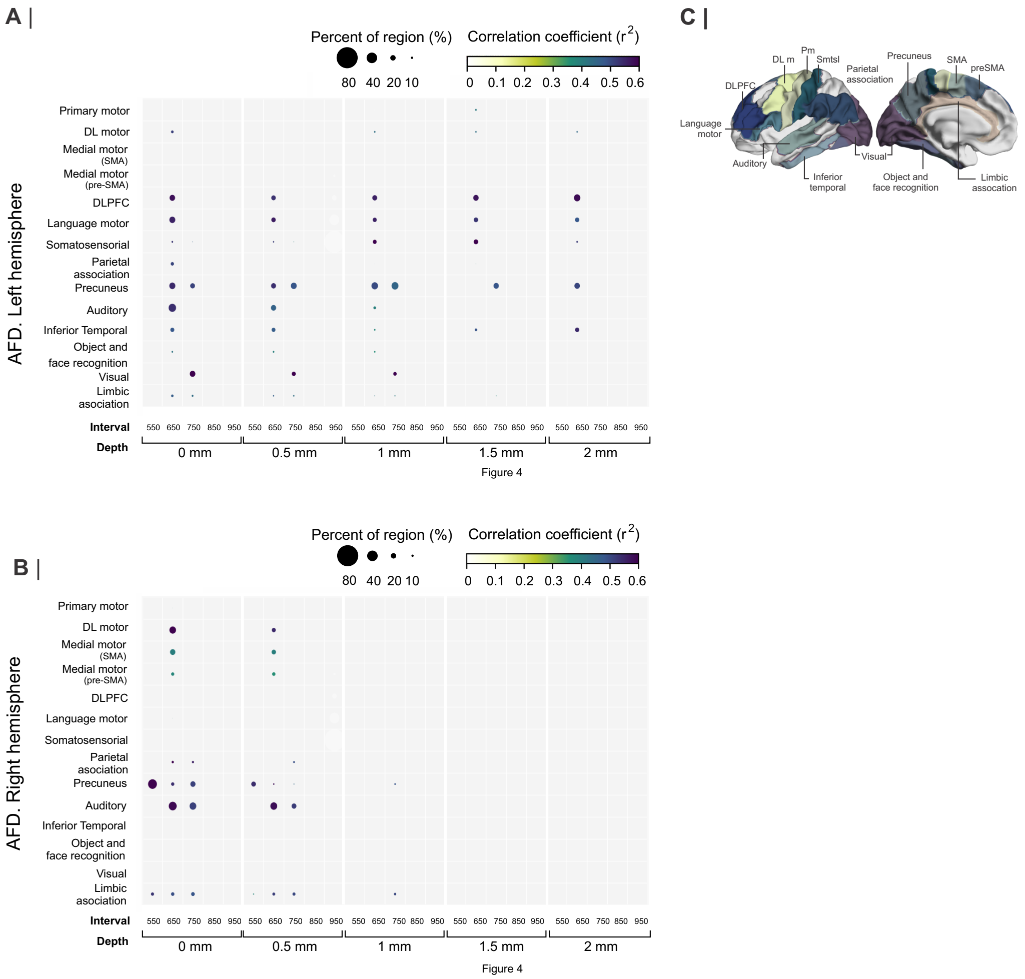


Extended data 7: A and B. Interregional correlation plot of the auditory asynchronies and $_{par}AFD$ as a function of the instructed interval and the depth of the $_{par}AFD$ calculation.

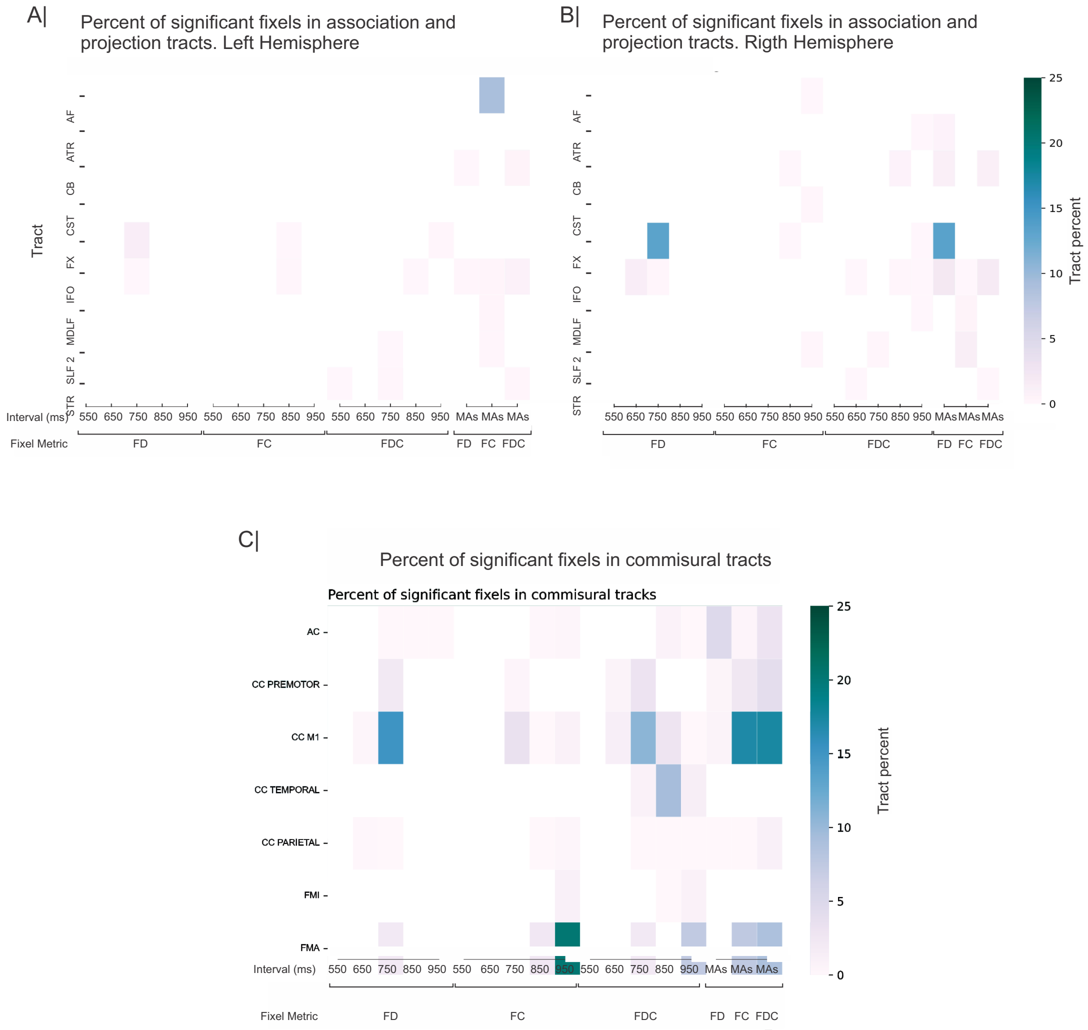


Extended data 8: Percent of significative fixels in the Deep White Matter in projection tracts (A) Left, (B) Right and in commissural tracts (C). AF= Arcuate fasciculus; ATR = Anterior Thalamic Radiation ; CB = cingulum bundle; CST= Corticospinal Tract ; FX = fornix; IFO= Inferior Longitudinal; MDLF= Middle Longitudinal Fascicle ; SLF 2= Superior Longitudinal Fasciculus 2; STR ; AC= Anterior commisure; CC = Corpus Callosum; FMI= Forceps Minior; FMA= Forceps Major; FD= Fiber Density; FC = Fiber Cross; FDC= Fiber Density and Cross; MAs = Mean Asychronies.
